# Shedding light on YfhS and YjlC: novel effectors of the NADH dehydrogenase activity of the electron transport chain in *Bacillus subtilis*

**DOI:** 10.64898/2026.03.25.714349

**Authors:** Carmella Gaucher, Logan Suits, Savannah Woods, Prahathees J. Eswara

**Affiliations:** Department of Molecular Biosciences, University of South Florida, Tampa, Florida, USA; Center for Antimicrobial Resistance, University of South Florida, Tampa, Florida, USA

**Author notes:** Address correspondence to Prahathees J. Eswara. Contributed equally (Carmella Gaucher, Logan Suits).

**Keywords:** Ndh, YjlC, NAD, NADH, ETC, YpsA, DUF1641

## Abstract

Oxidative phosphorylation is the most efficient way of generating ATP in respiring cells. As high energy electrons are the major source of reactive oxygen species their production needs to be carefully calibrated. In most organisms, NADH dehydrogenase serves as the primary source and gateway of electrons. This complex is responsible for oxidizing NADH to NAD^+^, which liberates two electrons that are then fed into the respiratory chain. In the Gram-positive model bacterium, *Bacillus subtilis*, a transcription factor (Rex) is utilized to monitor the rise in NADH level and subsequently increase the production of the NADH dehydrogenase Ndh. Thus, the generation of electrons through this pathway is tightly regulated. In this report, we reveal the presence of another independent mechanism to moderate Ndh activity involving a previously uncharacterized protein, YfhS. Additionally, we present the first experimental evidence showing that the functional NADH dehydrogenase is a two-protein complex comprised of a membrane-associated YjlC and the enzyme Ndh. We find that absence of YfhS leads to cell morphology and growth defects that are corrected by spontaneous mutations in *ndh*. We note that increased production of NADH dehydrogenase complex proteins by itself is not detrimental. However, strikingly, it is lethal in a strain lacking *yfhS*. These results reveal that YfhS is an important moderator of NADH dehydrogenase activity. We also demonstrate that YfhS and YjlC are interaction partners. A model developed based on our data indicates that YfhS is an important regulator of intracellular NADH concentration. Compounds that target specific microbial (Type II) NADH dehydrogenase, which is absent in human mitochondria, are considered promising drug candidates to help address the threat posed by antibiotic-resistant bacteria. Overall, our data unveiling the importance of YfhS and YjlC in controlling Ndh activity could be harnessed for the development of new therapeutics.

## INTRODUCTION

Cellular respiration is the most effective energy generator in nearly all life forms including bacteria, archaea, plants, and animals. In humans, defective mitochondrial respiratory complex is linked to numerous diseases (1, 2). During respiration, oxidation of high-energy electron donors such as NADH and FADH_2_ feed the electron transport chain (ETC) and synergistic export protons through the help of cytoplasmic membrane associated respiratory complexes. The proton gradient created by ETC subsequently powers the F_O_F_1_ ATP synthase to phosphorylate ADP thereby producing the energy-rich ATP. In aerobic organisms, the free electrons in ETC are used to reduce oxygen, the terminal electron acceptor, into water. In organisms that are capable of growing in the absence of oxygen, alternative terminal electron acceptors such as fumarate or nitrate are used to support anaerobic respiration. Additionally, certain organisms use less-productive substrate-level phosphorylation to generate ATP during fermentation. Although ETC is the most efficient way to produce energy, it is also the major source of reactive oxygen species (ROS) (3).

Redox enzymes, such as NADH dehydrogenases, are critical for energy generation. Three types of NADH dehydrogenases have been found in bacteria classified based on whether they pump protons (Type I; Nuo type; Complex I), sodium ions (Type III; Nqr), or neither (Type II) (4). The primary NADH dehydrogenase in the Gram-positive model organism *Bacillus subtilis* is the Type II enzyme Ndh (YjlD) (**Fig. 1A**) (5). To optimize electron transfer and minimize ROS generation (6-8), the production of redox enzymes needs to be tightly regulated. This sentinel function is performed by the transcription factor Rex which is responsible for assessing the levels of intracellular NADH. More specifically, high NADH concentration relieves Rex repression of *yjlC*-*ndh* operon and other genes involved in energetic pathways (**Fig. 1B**) (9-11). Consequently, Rex finetunes energy generation and metabolism in *B. subtilis* and other related organisms (11, 12). Availability of oxygen as a terminal electron acceptor and the presence of nutrients to generate NADH are intricately involved in ETC function. Therefore, transcriptional activity of *yjlC*-*ndh* operon is also under the control of the respiration-tuning ResDE two component system and stringent response (13, 14).

**Fig. 1:**
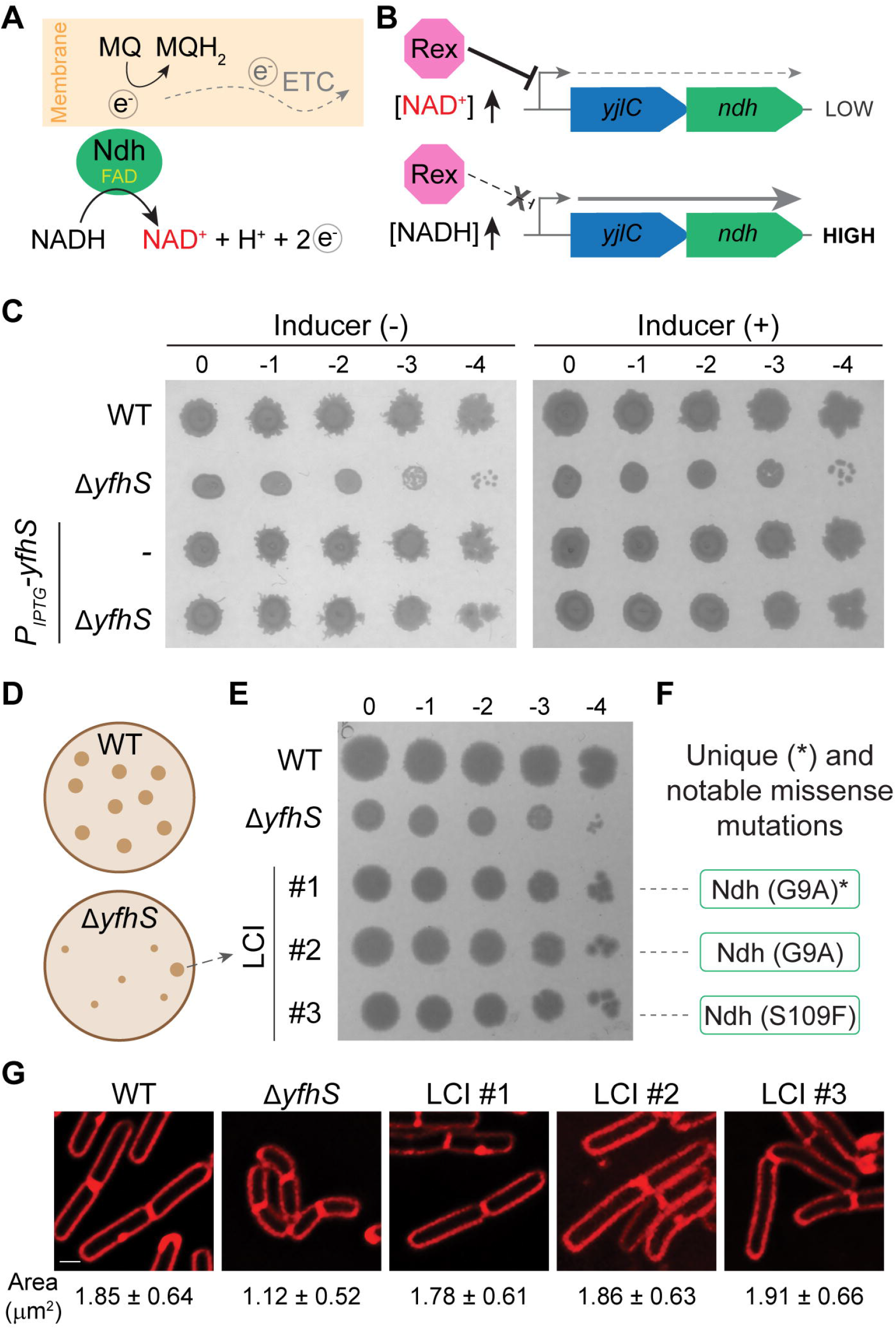
Spontaneous mutations in *ndh* suppress the small-colony and small-cell phenotypes of *yfhS* deletion mutant. **(A)** Illustration of electron transport chain (ETC) in *B. subtilis*. Ndh, bound to FAD cofactor, transfers electrons freed from the conversion NADH to NAD^+^ to reduce menaquinone-7 (MQ → MQH_2_), from which the electrons are eventually shuttled to the terminal electron acceptor typically oxygen. **(B)** Depiction of Rex mediated regulation of *yjlC*-*ndh* operon showing how the expression of these genes are differentially regulated based on NADH/NAD^+^ ratio. **(C)** Serial dilutions of WT (PY79), Δ*yfhS* (CG2), *yfhS*^*+*^ (CG172), and Δ*yfhS yfhS*^*+*^ (RB409). Representative pictures of plates incubated overnight at 37 °C are shown. Inducer plates contain 1 mM IPTG. **(D)** Illustration of plates showing colony size differences between WT and Δ*yfhS* strains. Spontaneous large colony forming mutants in Δ*yfhS* were isolated and screened as described in methods. **(E)** Representative plate with serial dilutions of WT (PY79), Δ*yfhS* (CG2), and three large colony isolates (LCIs; CG7, CG9, and CG10) incubated overnight at 37 °C is shown. **(F)** Suppressors found via whole genome sequencing. **(G)** Fluorescence micrographs of membrane stained (FM4-64, red) WT (PY79), Δ*yfhS* (CG2), and the three Δ*yfhS* LCIs (CG7, CG9, and CG10). For cross-sectional areas, n=100 per strain.

In a previous study from our laboratory, we discovered that *B. subtilis* cells lacking *yfhS* grow poorly and form small colonies on plate (**Fig. 1C**) (15). YfhS is a small protein (74 amino acids) of unknown function highly conserved in the Bacillales clade of Firmicutes phylum. Therefore, we investigated why the presence of YfhS is important for optimal growth rate. For this purpose, we capitalized on the observation that Δ*yfhS* strain spontaneously albeit less frequently formed large colonies on plate. Whole genome sequencing analysis of the stable large colony isolates revealed missense mutations in *ndh*. Thus, further experiments were designed to elucidate the relationship between Ndh and YfhS.

We note that deletion of both *rex* and *yfhS* leads to more severe growth inhibition. Additionally, we find that synthetic induction of *yjlC*-*ndh* (expression uncoupled from Rex control) is lethal in a strain lacking *yfhS*, but not in otherwise wild type (WT) or Δ*rex* backgrounds. This directly demonstrates a protective role of YfhS in moderating Ndh activity. Using luciferase-based *yjlC*-*ndh* promoter reporter we show that absence of *yfhS* leads to increased transcriptional activity suggesting elevated intracellular NADH concentration and Rex derepression. Similar increased activity was also seen when *yjlC* is knocked out. This result supports a recent prediction that YjlC, a DUF1641 domain containing protein, may serve as membrane anchor for Ndh to support its role in ETC (16). Furthermore, through bacterial two-hybrid analysis we detected direct interaction between YjlC-Ndh and YjlC-YfhS as well as self-interaction of all three proteins. In sum, our data demonstrates that YfhS is a key regulator of YjlC-Ndh NADH dehydrogenase complex. Fundamental studies such as ours are likely to inform translational research to address the growing threat of antibiotic resistance, as Type II NADH dehydrogenase is an attractive drug target (17-19).

## RESULTS

*yfhS* null phenotypes are corrected by spontaneous mutations in NADH dehydrogenase

In a previous study, we reported that *yfhS* null strain grows poorly compared to WT control (15). Using complementation analysis, we ensured that the absence of YfhS is specifically responsible for this growth defect as an inducible copy of *yfhS* reverses this phenotype even in the absence of inducer due to promoter leakiness (**Fig. 1C**) (15). In addition, we find that *yfhS* overexpression does not result in any observable phenotypic changes. While growing the Δ*yfhS* strain on a plate we mostly observed small colonies. Curiously, however few large colonies do spontaneously arise (**Fig. 1D**). We suspected that these large colonies held mechanistic clues regarding the physiological role of YfhS. Therefore, we set up an experiment to isolate large colony forming *yfhS* null mutants. After isolating and screening ten independent large colony isolates (LCI), we confirmed three that reproducibly formed large colonies (**Fig. 1E**). Additionally, we utilized fluorescence microscopy as another test to verify whether these LCIs correct Δ*yfhS* cell shape defects. As noted in our previous publication, *yfhS* null cells are smaller in comparison to WT (**Fig. 1G**) (15). This phenotype is corrected by *yfhS* complementation (**Fig. S2AB**). As cells harboring *yfhS* deletion are often curved, we quantified the cell surface area as depicted in **Fig. S2D**. The cell shape and area of Δ*yfhS* LCIs are more similar to WT than the Δ*yfhS* parent strain (**Fig. 1G**). Thus, both plate growth and cell morphology phenotypes of *yfhS* knockout mutant are corrected in LCIs. After ensuring this, we sequenced the whole genome of these LCIs to analyze what type of mutations are present and responsible for suppressing Δ*yfhS* phenotypes.

Sequencing results revealed that LCI #1 harbors a unique mutation in the *ndh* gene encoding NADH dehydrogenase which leads to a single amino acid change from Gly to Ala at position 9 (**Fig. 1F** and **Fig. S1**). Intriguingly, LCI #2 also has the same point mutation in *ndh* as LCI #1. However, this isolate also had an additional mutation in *ytxM* (Asp18Tyr). Intriguingly, this gene encodes an ETC-related protein YtxM/MenH which is involved in menaquinone biosynthesis (20, 21). Finally, LCI #3 also has a mutation in *ndh*, but in this case it is a Ser to Phe substitution at position 109. This strain also harbors an additional mutation in *ypzE* locus (G<u>C</u>T→G<u>A</u>T transversion mutation at genomic position 2228693). Although the role of *ypzE* is unclear it is found to be highly transcribed (22), and is possibly a 5’ untranslated region of the flavin mononucleotide riboswitch upstream of *ribU* gene which encodes a riboflavin transporter (23, 24). Notably flavin cofactors are often associated with redox enzymes of ETC (25). Ndh also uses flavin adenine dinucleotide as a cofactor. As *ndh* mutation is shared among all three LCIs, we tailored our experiments to decipher the functional relationship between YfhS and Ndh.

### YfhS is required to moderate Ndh activity

To test whether the LCI *ndh* mutations abolish the NADH dehydrogenase activity to restore the colony size of *yfhS* knockout strain, we generated a Δ*yfhS* Δ*ndh* double-deletion strain. As shown in **Fig. 2A**, deleting *ndh* by itself does not negatively influence the colony size. However, the colony size of Δ*yfhS* Δ*ndh* is similar to Δ*yfhS*. Thus, the LCI mutations are not rendering Ndh non-functional. Logically, we then wondered whether the LCI mutations increase the activity of Ndh. For this, we engineered strains to express *yjlC, ndh*, or *yjlC-ndh* under the control of an inducible promoter (*P*_*IPTG*_). The transcription from this promoter is therefore independent of Rex (**Fig. 1B**). We find that overexpression of *yjlC* or *ndh* in either WT or Δ*yfhS* backgrounds does not reveal anything significant as their colony morphologies resemble their corresponding parent strains (**Fig. 2B**). Astonishingly, we see that overexpression of *yjlC*-*ndh* in the absence of YfhS is lethal for cells, while that is not the case in WT background. This surprising result suggests that Ndh may require YjlC to form a functional NADH dehydrogenase complex. Overall, YfhS appears to be an important moderator of this complex (**Fig. 2C**).

**Fig. 2:**
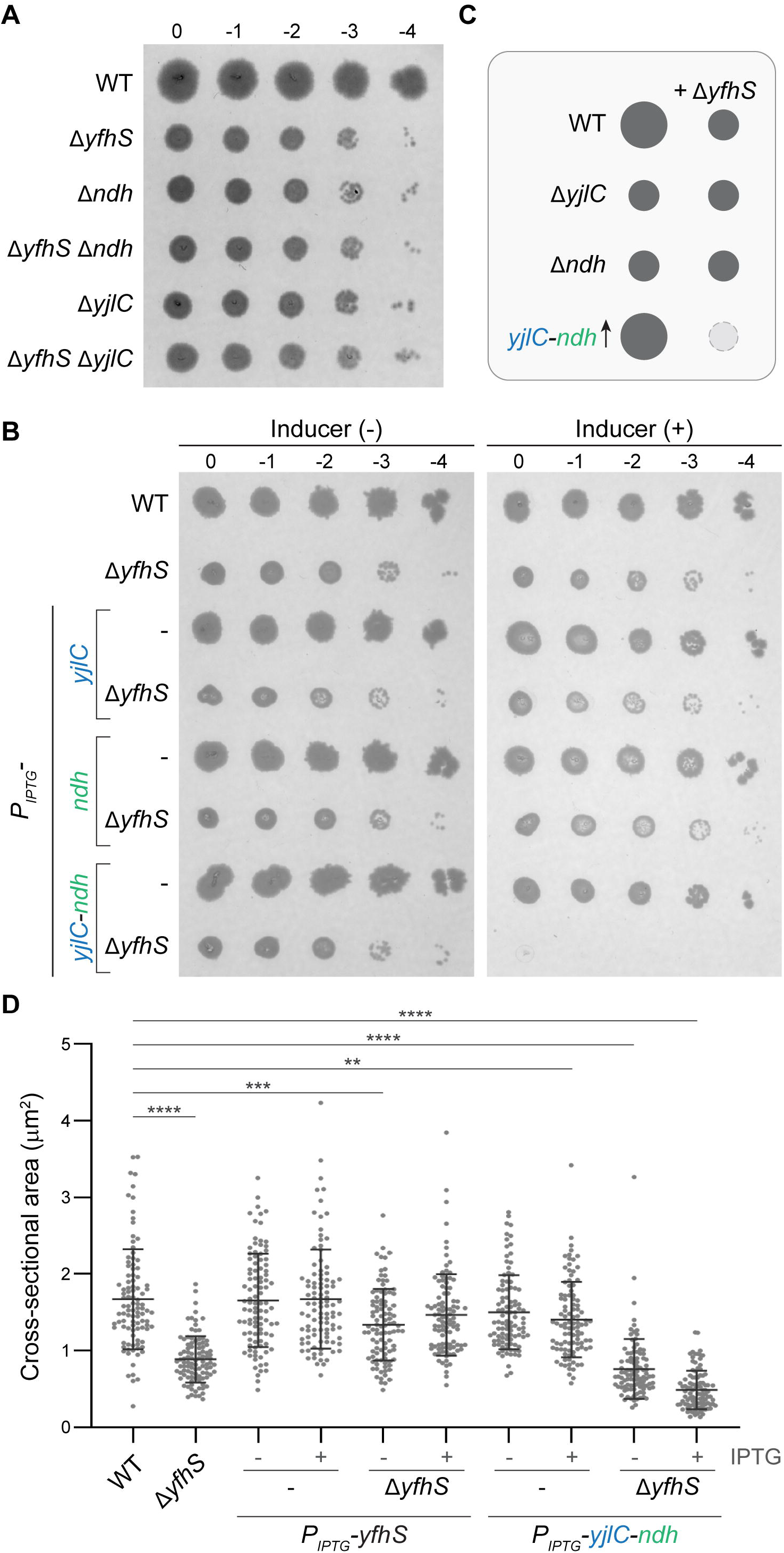
YfhS moderates YjlC-Ndh activity. **(A)** Serial dilutions of WT (PY79), Δ*yfhS* (CG2), Δ*ndh* (CG25), Δ*yfhS* Δ*ndh* (CG16), Δ*yjlC* (CG198), and Δ*yfhS* Δ*yjlC* (CG186). **(B)** Serial dilutions of WT (PY79), Δ*yfhS* (CG2), *yjlC*^*+*^ (CG135), Δ*yfhS yjlC*^*+*^ (CG136), *ndh*^*+*^ (CG143), and Δ*yfhS ndh*^*+*^ (CG144), and *yjlC-ndh*^*+*^ (BLS73), Δ*yfhS yjlC-ndh*^*+*^ (CG189). For both (A) and (B), representative pictures of plates incubated overnight at 37 °C are shown. Inducer plates contain 1 mM IPTG. **(C)** Illustration representing the effect of deleting *yjlC, ndh*, and *yfhS* on colony size. Also shown is the toxic phenotype of *yjlC-ndh* overexpression specifically in the absence of YfhS. **(D)** Cross-sectional areas of WT (PY79), Δ*yfhS* (CG2), *yfhS*^*+*^ (CG172), Δ*yfhS yfhS*^*+*^ (RB409), *yjlC-ndh*^*+*^ (BLS73), and Δ*yfhS yjlC-ndh*^*+*^ (CG189), grown in the presence of 1 mM IPTG where indicated. n=100 for all strains. ** = p<0.01, *** = p<0.001, **** = p<0.0001.

Next, we used high-resolution fluorescence microscopy to investigate the cause of lethality. As described earlier, the cross-sectional area of *yfhS* null mutant is significantly smaller than WT (**Fig. 2D**). We observed that overexpression of *yfhS* by itself does not exert any significant influence on the cell shape and that the characteristic Δ*yfhS* small-cell phenotype is complemented in the presence of inducible copy of *yfhS*. When *yjlC*-*ndh* is overexpressed in otherwise WT background, we find a modest but statistically significant reduction in the cell surface area. In contrast, we see that overproduction of YjlC and Ndh in the absence of YfhS leads to a stark decrease in cross-sectional area of cells (**Fig. 2D** and **Fig. S2CD**). In this strain, we would expect increased conversion of NADH to NAD^+^. Thus, the small-cell phenotype we observe is likely due to rapid depletion of NADH or accumulation of NAD^+^. In sum, our data establishes YfhS as a critical modulator of YjlC/Ndh activity.

### Absence of YfhS leads to increased transcription of *yjlC*-*ndh* operon

When intracellular NAD^+^ concentration is high, Rex represses the expression of *yjlC*-*ndh* operon (5, 11, 12). Conversely, increase in NADH level prevents Rex from being bound to DNA, and results in de-repression of this operon leading to increased production of YjlC and Ndh to help convert NADH to NAD^+^ (**Fig. 1B**). This allows for Rex-based finetuning of this operon simply based on the NADH/NAD^+^ ratio. This made us wonder whether the absence of Rex be detrimental in the absence of YfhS. We tested this by generating corresponding single and double deletion strains. As shown in **Fig. 3A**, while Δ*rex* colonies are similar to WT, we see pinprick colonies for the Δ*yfhS* Δ*rex* mutant that is different from the Δ*yfhS* strain (compare colonies indicated by arrowheads). Thus, either increased transcription of *yjlC*-*ndh* due to the absence of Rex or a subsequent potential drop in NADH concentration is detrimental to Δ*yfhS* cells. Next, we tested whether additional increase in YjlC/Ndh protein level is toxic in a Δ*rex* mutant using the IPTG-inducible *yjlC*-*ndh* construct. Unlike what we observe in *yfhS* null strain, overproduction of YjlC and Ndh is not detrimental in *rex* deletion mutant. Based on these results, we infer that the Rex-dependent transcriptional finetuning and more importantly YfhS-mediated regulation are needed to optimize the level and activity of YjlC/Ndh NADH dehydrogenase complex.

**Fig. 3:**
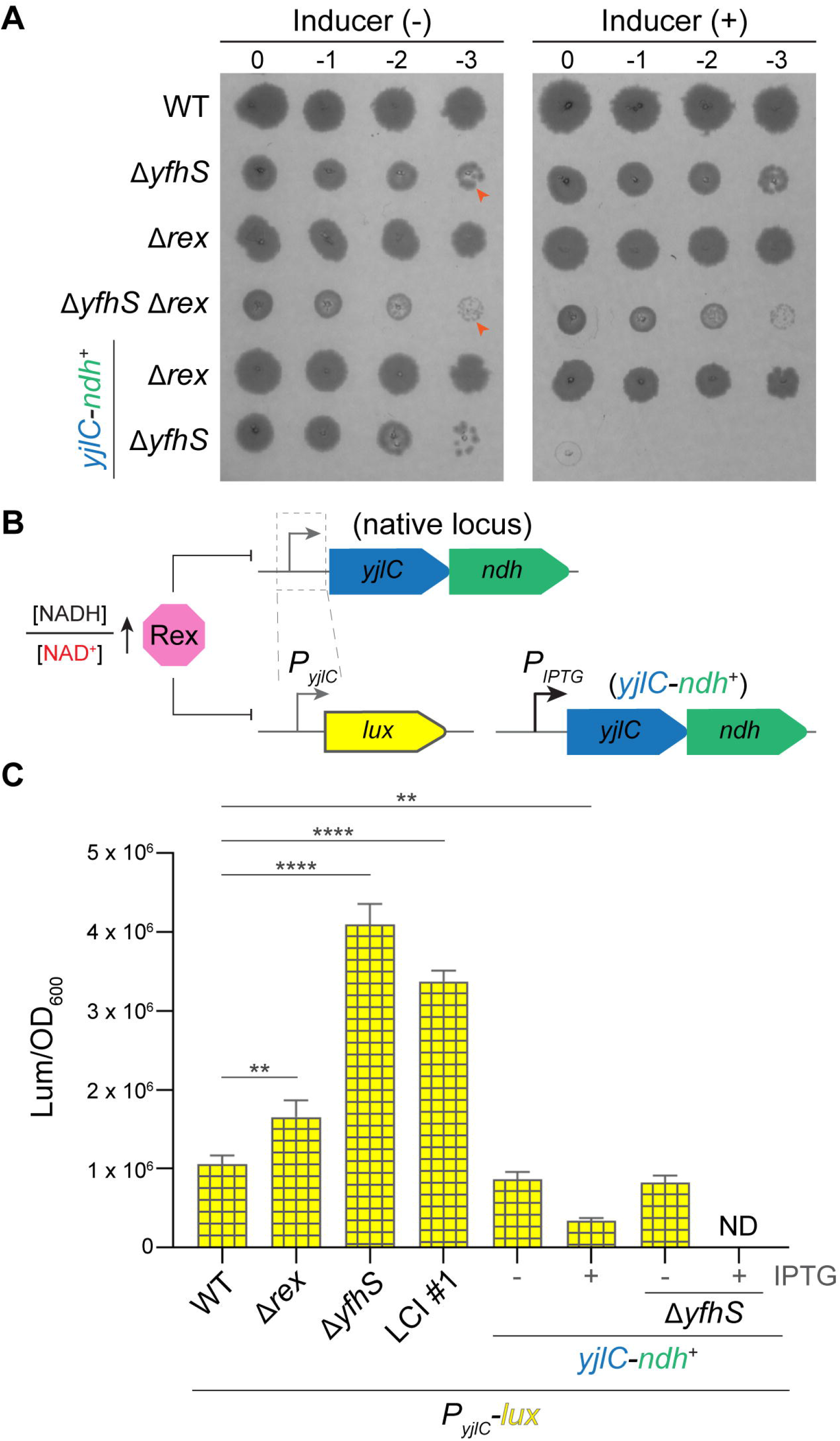
The promoter activity of *yjlC-ndh* operon increases in *yfhS* deletion strain. **(A)** Serial dilutions of WT (PY79), Δ*yfhS* (CG2), Δ*rex* (CG67), Δ*yfhS* Δ*rex* (CG68), Δ*rex yjlC*-*ndh*^+^ (CG69), and Δ*yfhS yjlC*-*ndh*^+^ (CG145). Arrowheads point to notable individual colony size differences. **(B)** Illustration showing the construction of luciferase (lux) based *P*_*yjlC*_*-lux* transcriptional reporter. In certain strains, *yjlC-ndh* operon is uncoupled from Rex control as well as other regulators of this promoters such as ResD with the help of an IPTG-inducible promoter at an ectopic locus. **(C)** Area under the curve of luminescence signal (arbitrary units) of *P*_*yjlC*_*-lux* reporter normalized by OD_600_ for WT (CG126), Δ*rex* (CG129), Δ*yfhS* (CG133), LCI #1 (CG130), *yjlC-ndh*^*+*^ (CG132), Δ*yfhS yjlC-ndh*^*+*^ (CG145). ND, not determined (due to toxicity).

Next, using a bacterial luciferase-based reporter (26), we monitored the transcriptional activity of *yjlC*-*ndh* promoter (*P*_*yjlC*_*-lux*) to assess the intracellular status of NADH/NAD^+^ ratio (**Fig. 3B**). As expected, we see increased promoter activity in Δ*rex* strain compared to WT (**Fig. 3C**). Interestingly, we see further elevation in *P*_*yjlC*_*-lux* transcription in the absence of YfhS. This heightened promoter activity compared to what we see in Δ*rex* implies the presence of additional factors, such as ResD and stringent response, that are linked to the transcription of *yjlC*-*ndh* operon (5, 13, 14). Intriguingly, we also observe higher promoter activity for LCI #1, a derivative of Δ*yfhS* parent which harbors a mutated *ndh* allele. This suggests that this Ndh (G9A) mutant is less efficient in generating NAD^+^, as it would be otherwise toxic (**Fig. 1E**). We also analyzed the promoter activity in WT and *yfhS* null strains overexpressing *yjlC*-*ndh*. Overexpression of this operon leads to decreased *P*_*yjlC*_*-lux* activity, hinting at a reduction in NADH levels. In contrast, when *yjlC*-*ndh* is overexpressed in Δ*yfhS* mutant background, we see severe growth defect and severely diminished luminescence (**Fig. S3**). Thus, our results indicate that the rapid depletion of intracellular NADH concentration is the likely reason to explain why YjlC/Ndh overproduction is lethal in cells lacking *yfhS* (**Fig. 3A**).

### YfhS-YjlC are interaction partners

We presumed that the YfhS-mediated modulation of NADH dehydrogenase activity may require direct protein-protein interaction. Therefore, we tested this possibility using bacterial two-hybrid analysis (27). Briefly, in this assay, negative or positive interaction between proteins is reported by colorless or blue color colonies respectively. We detect clear self-interaction for YjlC, Ndh, and YfhS (**Fig. 4A**). While Ndh self-association is known (4, 28), we provide the first evidence for YjlC and YfhS self-interaction to our knowledge. Additionally, we see a positive interaction between YjlC and Ndh which confirms a recent high-throughput proteomics-based observation (29) and a prediction based on modeling (16). Excitingly, we also see robust interaction between YfhS and YjlC. Thus, based on our results we can argue that YfhS directly interacts with YjlC to modulate Ndh activity.

**Fig. 4:**
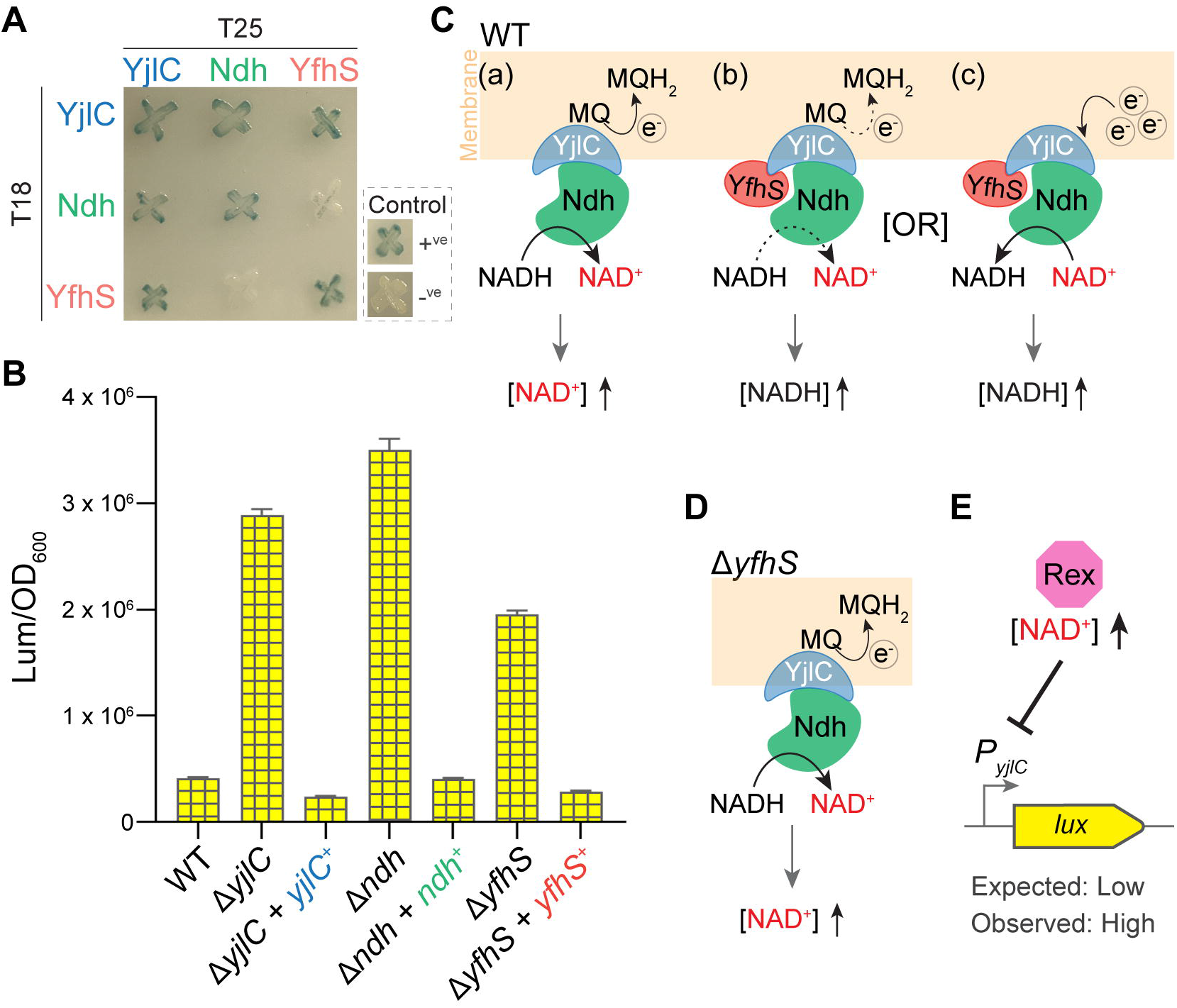
Protein-protein interaction analysis between YjlC, Ndh, YfhS. **(A)** Bacterial two-hybrid strains were grown on 5-bromo-4-chloro-3-indolyl-β-D-galactopyranoside (X-Gal) containing plate. Briefly, blue coloration indicates a positive interaction. YjlC, Ndh, and YfhS were probed for interaction with self and other partners in different combinations. **(B)** Activity of the *P*_*yjlC*_*-lux* reporter was assessed in WT (CG126), Δ*yjlC* (CG198), Δ*yjlC yjlC*^*+*^ (CG207), Δ*ndh* (CG199), Δ*ndh ndh*^*+*^ (CG208), Δ*yfhS* (CG200), Δ*yfhS yfhS*^*+*^ (CG202). **(C)** Possible models based on our results: YjlC and Ndh together form a functional NADH dehydrogenase complex (a); YfhS interaction with YjlC either decreases the rate of NADH to NAD^+^ conversion (b) or facilitates reverse electron transfer to generate NADH from NAD^+^ (c). **(D)** Based on the models discussed above (Cb and Cc), without YfhS, intracellular NAD^+^ level should be high. **(E)** Increased *P*_*yjlC*_ activity in Δ*yfhS* (in panel B) suggests that NADH level is higher (and not NAD^+^) similar to *yjlC* and *ndh* single deletion mutants. Thus, there is a contradiction between expected and observed results in Δ*yfhS* background. This discrepancy is likely due to the presence of additional Rex independent regulators of *yjlC*-*ndh* promoter such as ResD.

Apart from the clues presented in this study and the recent prediction (16), the essentiality of YjlC to support Ndh activity has not been experimentally validated. Therefore, we used our *P*_*yjlC*_*-lux* reporter to clarify this. As shown in **Fig. 4B**, we find that deletion of *yjlC* or *ndh* (also *yfhS* as shown previously) individually results in enhanced luminescence (high NADH and low NAD^+^) and this effect is reversed by the complementation of corresponding genes. We believe that this interaction and luminescence datasets, similar colony size of individual *yjlC*/*ndh* knockouts (**Fig. 2A**), as well as the toxicity we observe in Δ*yfhS* requiring overproduction of both YjlC and Ndh (**Fig. 2B**), establishes YjlC as an essential partner of the Type II NADH dehydrogenase complex.

As we see a positive interaction between YjlC-Ndh and YjlC-YfhS, we could speculate the following possibilities. **(i)** YfhS blocks the interaction between YjlC and Ndh which impairs conversion of NADH to NAD^+^. If this is true, then overproduction of YfhS would be expected to lead to NADH accumulation (increased *P*_*yjlC*_*-lux* promoter activity). However, this is not what we observe (**Fig. 4B**). In fact, we see the opposite. Thus, we do not think YfhS obstructs YjlC-Ndh interaction. **(ii)** YfhS-YjlC interaction is needed for Ndh to function. In this case, deletion of *yfhS* or *yjlC* should exhibit similar extent of elevated promoter activity. However, the signal is highly elevated for Δ*yjlC* and Δ*ndh* but not to the same degree for Δ*yfhS* (**Fig. 4B**). Additionally, if this was true then overexpression of *yjlC*-*ndh* in the absence of YfhS should not be toxic (**Fig. 2B**). However, our observation contradicts this prediction. As such, we do not believe that YfhS is required for the function of YjlC/Ndh complex to generate NAD^+^ (**Fig. 4Ca**). **(iii)** YfhS-YjlC interaction negatively modulates the NADH dehydrogenase activity. Perhaps the presence of YfhS within the complex dampens the Ndh activity (**Fig. 4Cb**). So, deletion of *rex* or overexpression of *yjlC*-*ndh* does not exhibit any detrimental effect while YfhS is present (**Figs. 2B** and **3A**). **(iv)** Alternatively, YfhS somehow promotes reverse electron transfer (RET) to generate NADH from NAD^+^ (**Fig. 4Cc**) (30). Thus, WT cells are able to balance the NADH/NAD^+^ ratio as required. However, in either cases (iii) or (iv), deletion of *yfhS* should lead to increased intracellular NAD^+^ level (**Fig. 4D**). If so, we would expect *P*_*yjlC*_*-lux* activity to be low, not high (**Fig. 4E**). Accordingly, the discrepancy in our model indicates the involvement of another factor that is also presumably regulating this promoter. A well-documented regulator of *yjlC*-*ndh* operon is the oxygen-responsive ResDE two-component system (13, 31-33). Specifically, it was found that *resD* deletion leads to upregulation of this operon in a manner independent of Rex (33). Therefore, we investigated the potential relationship between YfhS and ResDE.

### YfhS may be involved in ResDE signaling

Using our *P*_*yjlC*_*-lux* reporter, we first investigated the luminescence produced in *resD* knockout strain. As expected, we observed enhanced signal greater than *yfhS* and *rex* individual deletion strains (**Fig. 5A**). Double deletion of *rex* and *resD* nearly resembled Δ*resD*. Thus, ResD repression of *yjlC*-*ndh* promoter appears to be stronger than that of Rex. Upon closer inspection, we find that deletion of *rex* alone results in elevated promoter activity only during the exponential phase (**Fig. S4**). Conversely, ResD appears to strongly influence the expression of this operon during post-exponential phase as *P*_*yjlC*_*-lux* activity in Δ*resD* appears to be the inverse of Δ*rex* during this growth period. Deletion of both *resD* and *rex* leads to significantly heightened promoter activity throughout nearly all growth phases. Based on these results, we could hypothesize that YfhS collaborates with YjlC to communicate the status of NADH dehydrogenase activity or its accompanying electron flow, to toggle ResE sensor protein between its kinase and phosphatase states either directly or indirectly (**Fig. S5**) (34-36). In this scenario, ResE is kept in a kinase “off”/phosphatase “on” state by YfhS when Ndh functions, presumably in the presence of oxygen as the terminal electron acceptor (**Fig. 5B** and **Fig. S5A**). If this speculation is true, then in the absence of YfhS, ResE will be kinase “on”/phosphatase “off” intermittently, and as a result ResD will be unable to strongly repress the expression of the *yjlC*-*ndh* operon (**Fig. S5B**). If the poor growth and increased *P*_*yjlC*_*-lux* activity we observe in Δ*yfhS* is due to partial ResD de-repression, then *resE* deletion should reverse this effect (**Fig. S5C**). We tested this prediction by analyzing the colony morphologies of all relevant strains on a spot titer plate. As shown in **Fig. 5C**, our results confirm that double-deletion of *resE* and *yfhS* does indeed allow cells to grow similar to WT. Additionally, we note that the strain harboring *resD* deletion exhibits small colony phenotype comparable to Δ*yfhS*. Together these results indicate that YfhS modulates the intracellular NADH/NAD^+^ ratio through ResDE two component system. Whether this is achieved directly or with the help of other factor(s) remains to be explored.

**Fig. 5:**
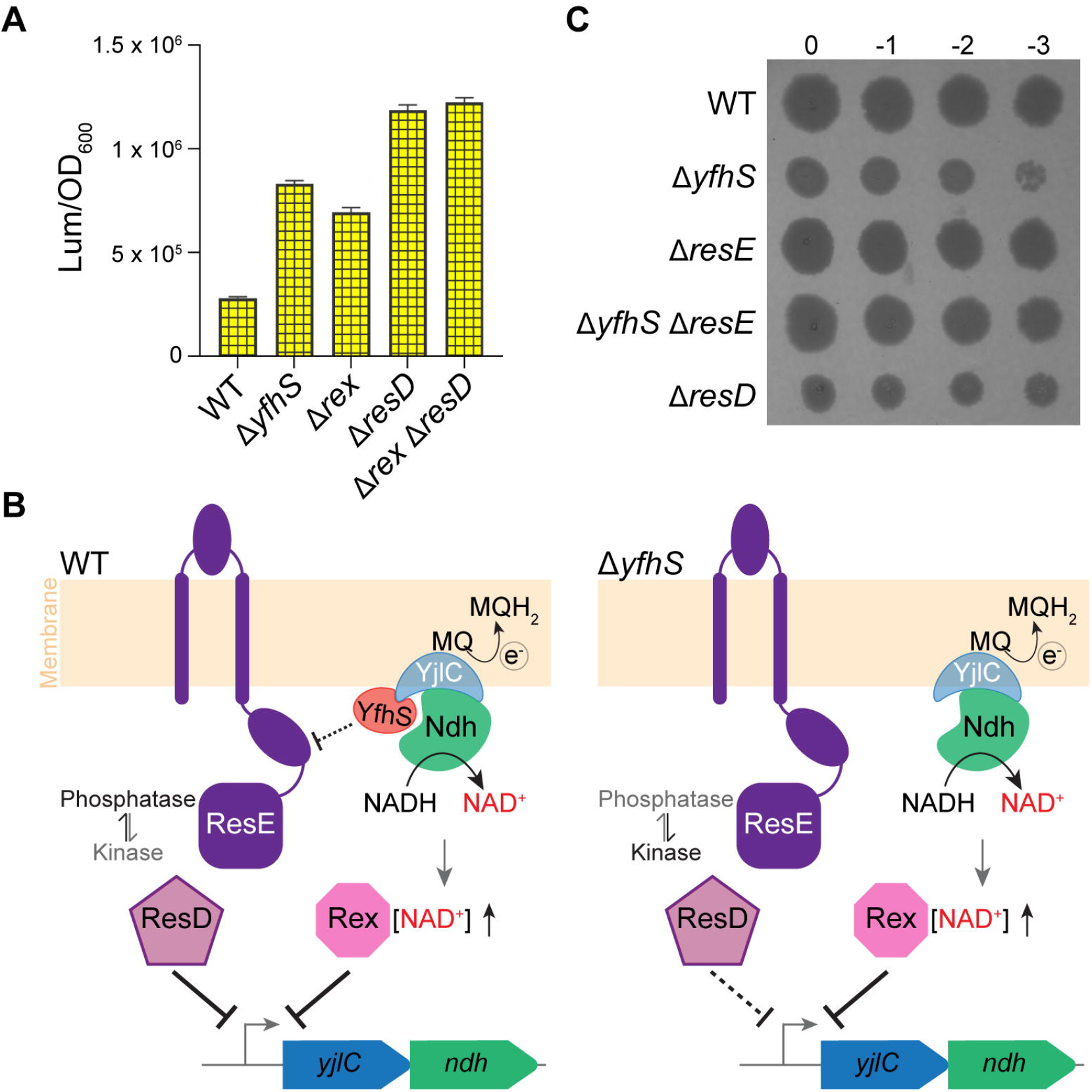
Deletion of *resE* reverses the Δ*yfhS* growth phenotype. **(A)** Area under the curve for luminescence divided by OD_600_ for WT (CG126), Δ*yfhS* (CG194), Δ*rex* (CG129), Δ*resD* (CG218), Δ*rex* Δ*resD* (CG213). **(B)** Working model to illustrate the putative role of YfhS in ResDE signaling. When present in complex with YjlC, YfhS may either directly or indirectly inhibit the kinase activity of ResE and favor the phosphatase activity, thus allowing ResD mediated strong repression of *P*_*yjlC*_. Without YfhS, the kinase/phosphatase functions of ResE may not be tunable. As such, we would expect weak repression (dotted line) of *yjlC*-*ndh* promoter by ResD, leading to elevated *P*_*yjlC*_ activity that we observe in Δ*yfhS* (for more detailed models, see Fig. S5). **(C)** Serial dilutions of WT (PY79), Δ*yfhS* (CG2), Δ*resE* (CG221), Δ*yfhS* Δ*resE* (CG220), and Δ*resD* (CG212).

## DISCUSSION

We first noted the cell and colony morphology phenotypes of Δ*yfhS* strain while investigating *yfhS* disruption which suppressed the toxic phenotype associated with another protein of unknown function YpsA (15). Interestingly, YpsA belongs to the SLOG superfamily of proteins predicted to bind nucleotides similar to NAD^+^ (37, 38). In light of the results provided in this article, perhaps absence of YfhS possibly alleviates the toxicity of YpsA overproduction by altering the intracellular concentration of NAD^+^.

In this report, we have shown that cells lacking YfhS exhibit defective cell shape and growth phenotypes. By studying the spontaneous large colony forming *yfhS* null mutants, we discovered that YfhS plays a crucial role in regulating YjlC-Ndh activity. We see enhanced luminescence using our *P*_*yjlC*_*-lux* reporter in the absence of YfhS indicating altered NADH/NAD^+^ ratio. Interestingly, the pattern of *P*_*yjlC*_*-lux* expression is distinct between WT and *rex* deletion strains only during exponential phase, where the latter shows elevated promoter activity (**Fig. S3**). In contrast, the promoter activity of *yfhS* null strain and LCI are elevated in the post-exponential phase. Thus, as discussed below, the increased luminescence we see in Δ*yfhS* mutant backgrounds is likely due to impaired ResD repression (**Fig. S5AB**) (5, 13). Regardless, as overexpression of *yjlC*-*ndh* is toxic to Δ*yfhS* cells we can argue that the small-cell and poor growth phenotypes are likely due to hyperactive YjlC-Ndh complex. Consequently, LCIs correct this defect by mutating *ndh*. Both Ndh mutations we noted in our LCIs (G9A and S109F) are located in the FAD cofactor binding pocket (**Fig. S1**) (4). As such, although highly expressed (**Fig. 3C**), we suspect that these mutants are less efficient and help maintain normal NADH levels needed to support growth.

Proper NADH/NAD^+^ ratio is essential for growth rate optimization, as we see small colonies for individual deletions of *yjlC, ndh, yfhS*, and *resD* (**Fig. 2A** and **Fig. 5C**). While absence of YjlC or Ndh would lead to accumulation of NADH, we think lack of YfhS or ResD is possibly elevating NAD^+^ concentration in the cells. We see that cells lacking *yfhS* are unable to tolerate increased production of YjlC and Ndh (**Fig. 2B** and **Fig. 3A**). In this case, we observe decreased *P*_*yjlC*_*-lux* promoter activity (**Fig. 3C** and **Fig. S3**) suggesting that intracellular NAD^+^ concentration is possibly high in this strain. Therefore, we believe that rapid depletion of NADH is the cause of toxicity in this strain (**Fig. S5F**). As *resE yfhS* double deletion strain forms normal size colony (**Fig. 5C**), we infer that at least partial repression of *yjlC*-*ndh* operon by ResD alone is sufficient for optimizing the NADH/NAD^+^ ratio presumably through Rex (**Fig. S5C**). Nevertheless, why overexpression of *yjlC*-*ndh*, uncoupled from Rex and ResD control, is not toxic by itself but only when YfhS is absent is puzzling (**Fig. 2B**). If YfhS is capable of allowing RET (**Fig. 4Cc**), then that would potentially address this conundrum, as illustrated in **Fig. S5FG**. According to our speculation, in wild type cells, YfhS may assist in RET specifically when there is electron overflow. This would simply allow the cells to maintain the levels of NADH and NAD+ in the normal range even without the help of ResD and Rex transcription factors. If our model is true, then how overflow of electrons is sensed by YfhS would need clarification in the future. Our model also explains why we see moderate and severe small colony phenotypes for Δ*yfhS* and Δ*yfhS* Δ*rex* respectively, but not Δ*rex* (**Fig. S5D-E**). Interestingly, the expression of *yfhS* is found to be highly elevated in cells undergoing anaerobic growth or sporulation (**Fig. S6A**) (39, 40). Thus, it is tempting to speculate that YfhS signaling through ResE pathway and/or RET mediated production of NADH is especially helpful during these conditions. Additionally, sporulation is triggered in response to nutrient starvation (41), a condition that initiates stringent response (42). Intriguingly, *yjlC*-*ndh* expression is negatively regulated by stringent response (14). Therefore, the transcription profile of *yfhS* and *yjlC*-*ndh* are inversely correlated specifically during anoxic period and nutrient limiting conditions that activate sporulation (**Fig. S6A**) (39, 40). Hence, it is tempting to speculate that generation of NADH from NAD^+^ through RET may be favored during these specific circumstances to maintain the redox buffering capacity required for survival or spore maturation (43). However, further experiments are warranted to specifically test whether YfhS participates in or promotes RET.

Phylogenetic gene conservation and genetic neighborhood analysis using GeCoViz (44), revealed that *yfhS* is highly conserved in the Firmicutes phylum specifically within the Bacillales order which includes all the aerobic endospore-forming bacteria (**Fig. S7**). Thus, it is possible the presence of YfhS assists in the process of sporulation specifically in oxic environment. We see strong synteny for *yfhS* with *mutY, fabL*, and *yfhP*. In *B. subtilis*, YfhP appears to be the transcription factor responsible for negatively regulating the expression of *mutY* and *fabL* genes (**Fig. S6B**) (45). Interestingly, MutY is a DNA repair enzyme involved in repairing DNA lesions (8-oxoG) caused by oxidative stress using oxidative stress-responsive iron-sulfur cluster as co-factor (46-48). FabL is a NADPH consuming oxidoreductase involved in lipid biogenesis. It has been shown that FabL is synthetically essential with FabI which consumes NADH (49). Thus, *B. subtilis* cells are equipped to continue the essential process of fatty acid synthesis when NADH is lacking by using NADPH. Given the role of YfhS in maintaining NADH/NAD^+^ ratio and preventing oxidative stress stemming from dysfunctional NADH dehydrogenase, the genomic position of *yfhS* in *B. subtilis* and other organisms, encoded on the opposite strand sandwiched between *mutY* and *fabL* is highly intriguing (**Fig. S6B** and **Fig. S7**). Similar analysis for *yjlC* uncovered that it is widely conserved in the kingdoms on both Bacteria and Archaea (**Fig. S8**), as noted in a recent article (16). Synteny between *yjlC* and genes encoding proteins with putative FAD binding domain and NADH oxidation or formate dehydrogenase function is easily recognizable.

Overall, in this study, we have elucidated the role of YfhS and YjlC as crucial regulators of the Type II NADH dehydrogenase activity. Thus, their respective functions help cells maintain a healthy intracellular NADH/NAD^+^ ratio. *B. subtilis* is used in biotechnology industry where pathways involving NADH are altered to increase the yield of metabolic byproducts (50, 51). Therefore, identification of new players involved in this pathway would likely assist in increasing the metabolite yield. More importantly, absence of Type II NADH dehydrogenase in humans underscores the potential of the new proteins we have characterized in this study, YfhS and YjlC, as attractive drug targets (17-19).

## MATERIALS AND METHODS

### Strain construction

All *B. subtilis* strains used in this study are derivatives of PY79 (52). Information pertaining to strain construction is provided in the supplemental file. Strains and oligonucleotides used in this report are listed in **Table S1** and **Table S2** respectively. The bacterial two-hybrid assay related *E. coli* strains and plasmids are also listed in the supplemental tables (27). *B. subtilis* non-essential gene knockout mutants (53), the plasmid for removing the resistance marker cassette (pDR244), and the vectors for generating luciferase reporter fusions (pBS3Klux/pBS3Elux) (26) were obtained from the Bacillus Genetic Stock Center. Plasmid pDR111 (David Rudner) was used for generating IPTG-inducible constructs. Each plasmid was confirmed by Sanger sequencing or whole plasmid sequencing (Azenta/Genewiz). These newly transformed strains were confirmed through selective plating and PCR.

### Culture media

All experiments were conducted in lysogeny broth (LB) for liquid culture assays and lysogeny broth agar (LA) for plate assays.

### Spot titer assay

Cell grown overnight in LB at 30 °C were back-diluted to an OD_600_ of 0.1. The mid-exponential phase cells post 2 h incubation at 37 °C, were 10-fold serially diluted and 1 µl of each dilutions were spotted on LA (with or without inducer) using a multichannel pipette. Plates were imaged after overnight incubation at 37 °C.

### Isolation of large colony mutants

The Δ*yfhS* strain (CG2) from glycerol stock was streaked out on multiple LA plates for single colony isolation and incubated at 37 °C overnight. An individual colony from each plate was used for inoculation and grown in LB for 6 h. Mid-exponential phase cultures were serially diluted, and 10^-6^-dilution was plated on LA using sterile glass beads and incubated overnight at 37 °C. The following day, if a large colony appeared among the otherwise small colonies (**Fig. 1D**), one colony per plate was then selected and re-streaked for further isolation and screening.

### Microscopy

Mid-exponential phase cells were imaged. For experiments requiring addition of an inducer, cells were grown in the presence of IPTG for 2 h prior to imaging. The cells were then stained with SynaptoRed (FM4-64; 1 μg/ml) membrane dye. Sample preparation was carried out as per our published protocol (54). Images were acquired using Zeiss Axio Observer 7 microscope. ImageJ was used to quantify the cross-sectional area of the cells and to perform statistical analysis.

### Luminescence assay

The strains harboring *P*_*yjlC*_*-lux* reporter were grown overnight in LB at 30 °C. The overnight cultures were then back-diluted to an OD_600_ of 0.1 in 1 mL LB with or without inducer as needed. The cultures were then grown at 37 °C shaking incubator in 15 mL falcon tubes for 1 h. Following this step, 100 μl of each strain was pipetted into a 96-well plate and placed in a microplate reader (BioTek Synergy H1) at 37 °C with shaking enabled. The luminescence and OD_600_ were monitored for over 16 hours. To analyze the data, the luminescence readings were averaged for every technical replicate and normalized by the averaged OD_600_ and graphed with GraphPad Prism. GraphPad was used to calculate the area under the curve (AUC) and for conducting standard statistical analysis. One-way Anova was applied to determine the statistical significance.

### Bacterial two-hybrid assay

The bacterial two-hybrid plasmids harboring our genes of interest were transformed into BTH101 *E. coli* competent cells. The possibility of interaction between proteins were then probed using chromogenic plates as described previously (27, 55).

## ACKNOWLEDGEMENTS

We thank the Eswara laboratory members for their helpful comments and productive discussions on this project. This work was funded by the National Institutes of Health (R35GM133617 to PJE) and the University of South Florida - Center for Antimicrobial Resistance award (PJE). The funders had no role in study design, data collection and analysis, decision to publish, or preparation of the manuscript.

## AUTHOR CONTRIBUTIONS

Study design (CG, LS, and PJE), strain construction and data acquisition (CG, LS, SW, and PJE), data analysis (CG, LS, SW, and PJE), writing of the manuscript (CG, LS, SW, and PJE), resources (PJE), funding (PJE), and supervision (PJE).

