## Supplemental Information for "Shedding light on YfhS and YjlC: novel effectors of the NADH dehydrogenase activity of the electron transport chain in *Bacillus subtilis*"

### SUPPLEMENTAL FIGURE AND TABLE LEGENDS

**Supplemental figure 1: Multiple sequence alignment of Ndh homologs.** Sequences of *Bacillus subtilis* (*Bs*), *Caldalkalibacillus thermarum* (*Cth*), *Staphylococcus aureus* (*Sa*), and *Mycobacterium tuberculosis* (*Mtb*) Type II NADH dehydrogenases were aligned using Clustal Omega. Residues highlighted in yellow are part of the FAD binding pocket as determined from the *Cth* Ndh crystal structure. GXGXXG and AQXAXQ are putative nucleotide and quinone binding motifs respectively. Suppressor mutations identified in this study (G9A and S109F) are shown in red font and underlined.

**Supplemental figure 2: Analysis of small-cell phenotypes associated with *yfhS* deletion.** **(A)** Micrographs of WT (PY79) and Δ*yfhS* (CG2). **(B)** Micrographs of *yfhS*$^+$ (CG172) and Δ*yfhS yfhS*$^+$ (RB409). **(C)** Micrographs of *yjlC-ndh*$^+$ (BLS73) and Δ*yfhS yjlC-ndh*$^+$ (CG189). All strains were grown in LB without (-) or with (+) 1 mM IPTG. Cells were stained with FM4-64 membrane stain (red). Scale bar, 1 µm. **(D)** Quantification and illustration of 2-dimensional cross-sectional area measurements of WT (PY79), Δ*yfhS* (CG2), and Δ*yfhS yjlC-ndh*$^+$ (CG189) grown in the presence of inducer. The means and standard deviations are shown, n=100.

**Supplementary figure 3: Linear curves of data shown in Fig. 3C.** Luminescence (top panel; arbitrary units) and $OD_{600}$ (bottom panel) of WT (CG126), Δ*rex* (CG129), Δ*yfhS* (CG133), LCI #1 (CG7), *yjlC-ndh*$^+$ (CG132), and Δ*yfhS yjlC-ndh*$^+$ (CG189) grown without (-) or with (+) 1 mM IPTG. Note: IPTG addition is inhibitory to the growth of Δ*yfhS yjlC-ndh*$^+$ strain.

**Supplemental figure 4: Linear graph associated with the data shown in Fig. 5A.** Luminescence data normalized by $OD_{600}$ is shown for WT (CG126), Δ*rex* (CG129), Δ*yfhS* (CG133), Δ*resD* (CG218), and Δ*resD* Δ*rex* (CG213) strains.

**Supplemental figure 5: Hypothetical models to explain the various contradictory results.** **(A)** Wild type (WT) cells are able to finetune the NADH/$NAD^+$ ratio by efficiently utilizing both ResD and Rex transcription factors. The putative role of YfhS in promoting reverse electron transfer (RET) and influencing ResDE signaling is depicted in dotted lines. Specifically, we predict that the kinase/phosphatase functions of ResDE is altered by YfhS. **(B)** In the absence of YfhS, cells are unable to fine-tune ResDE signaling (weak repression from ResD) and rely more on Rex. This impaired repression of *yjlC*-*ndh* expression leads to excess conversion of NADH to $NAD^+$ and subsequent accumulation of electrons in the ETC system. Thus, either NADH depletion or excess electrons (and associated ROS stress) manifests as small-colony phenotype (panel G). **(C)** In the absence of both YfhS and ResE, ResD will be expected to repress the *yjlC*-*ndh* operon. In this case, cells prevent increased expression of this operon and also make use of Rex mediated regulation. This minimizes the detrimental effects of NADH consumption. As such, the colony size of this strain resembles WT (see panel G). **(D)** Cells lacking *rex* are not severely affected as proper NADH/$NAD^+$ ratio is maintained by YfhS through ResDE signaling (refer to panel G). **(E)** Deletion of both *rex* and *yfhS* is harmful. In this scenario: (i) cells are unable to utilize the ResDE pathway properly as explained in (B); and (ii) absence of Rex prevents the secondary gate-keeping mechanism to regulate the expression of *yjlC*-*ndh* operon. This leads to heightened consumption of NADH and increased production of high-energy electrons which leads to poor growth as depicted in panel G. **(F)** We were baffled by the observation that uncoupling the expression of *yjlC*-*ndh* operon from ResD and Rex control (using an IPTG-inducible promoter; *yjlC*-*ndh* ↑) is harmless to otherwise WT strain but is lethal when YfhS is absent. This result suggested that the presence of YfhS is more critical than the ResD/Rex transcription factors. We suspect that YfhS-YjlC interaction (Fig. 4A) is key to this protective effect. According to our speculative model, YfhS serves similar to an “electrostatic discharge device” to safely neutralize electron buildup by facilitating RET. In the absence of such a mechanism in the Δ*yfhS* strain background, cells are unable to properly ground themselves. Thus, it is lethal to uncouple the expression of *yjlC*-*ndh* from ResD and Rex, specifically in the absence of YfhS (see panel G). **(G)** Pictographic summary of colony morphologies observed for various strains discussed in this figure. NG, no growth.

**Supplemental figure 6: Gene expression profiles of *yfhS*, *yjlC*, and *ndh* and gene neighborhood of *yfhS*. (A)** Transcriptomics data showing that the expression pattern of *yfhS* and *yjlC-ndh* operon is inversely correlated during anaerobic growth and sporulation. **(B)** *yfhS* is encoded within the *mutY* and *fabL* locus on the complementary strand.

**Supplemental figure 7: *yfhS* is conserved within Bacillales.** Analysis using GeCoViz reveals that *yfhS* is highly conserved within the Bacillales order of the Firmicutes phylum. Strong synteny is noted with a gene involved in repairing DNA lesions (8-oxoG) caused during oxidative stress (MutY) and another gene that encodes a protein with putative oxidoreductase function (in *B. subtilis* it is FabL).

**Supplemental figure 8: Inter-kingdom conservation of *yjlC*.** GeCoViz analysis uncovered that *yjlC*-like genes are broadly conserved in organisms that belong to the kingdom of Bacteria as well as Archaea. Clear synteny exists with genes that encode proteins with FAD binding domain, NADH oxidation, or formate dehydrogenase function.

**Supplemental Table 1: Strains used in this study**

**Supplemental Table 2: Oligonucleotide primers used in this study**

### SUPPLEMENTAL METHODS

#### Plasmid Construction

*B. subtilis* IPTG-inducible gene constructs:

*yfhS* overexpression plasmid: pRB54

To create the plasmid pRB54, *yfhS* was amplified from PY79 Chromosomal (Ch) DNA using primers oRB59/oRB60. This PCR product was then digested with the enzymes SalI and NheI and ligated into pDR111 vector also cut with SalI/NheI.

*yjlC* overexpression plasmid: pLS27

To create the plasmid pLS27, *yjlC* was amplified from PY79 Ch DNA using the primer pairs oLS41/oLS42. This fragment was cloned during SalI/NheI restriction enzymes and ligated into a pDR111 vector.

*ndh* overexpression plasmid: pCG23

The plasmid pCG23 was created using primer pairs oLS43/oCG59 to amplify *ndh* from PY79 Ch DNA. Traditional cloning approach involving SalI/NheI cut sites were used to place *ndh* into pDR111.

*yjlC-ndh* overexpression plasmid: pLS28

To create the plasmid pLS28, we amplified *yjlC-ndh* region from the PY79 DNA using primer pairs oLS41/oLS44 and cloned it into pDR111 using SalI/NheI cut sites.

Gene expression monitoring using luciferase:

$P_{yjlC}$-*lux* transcriptional reporter: pCG22

The primer pairs oCG29/oCG30 were used to linearize the empty vector (pBS3Klux). The primer pairs oCG31/oCG32 were used to amplify the promoter of the *yjlC-ndh* operon (485 bp region upstream of start codon). Gibson Assembly was used to stitch the fragments together.

### Table S1: Strains used in this study

(*BKE/BKK strains were obtained from BGSC)

| Strain | Genotype | Reference |
|---|---|---|
| PY79 | *B. subtilis* (wild type) | Youngman et al., 1984 |
| CG2 | *yfhS::erm* | *BKE08640 → PY79 |
| AHB286 | *bkdB::Tn917::amyE::cat* | Amy Camp |
| CG172 | *bkdB::Tn917::amyE::*$P_{hyspank}$*-yfhS spec* | pRB54 → AHB286 |
| RB409 | Δ*yfhS::erm bkdB::Tn917::amyE::*$P_{hyspank}$*-yfhS spec* | *BKE0840 → AHB286 |
| CG7 | Δ*yfhS::erm ndh* suppressor *ndh* G9A (LCI #1) | CG2 → PY79 |
| CG9 | Δ*yfhS::erm ndh* suppressor (LCI #2) | CG2 → PY79 |
| CG10 | Δ*yfhS::erm ndh* suppressor (LCI #3) | CG2 → PY79 |
| CG25 | *ndh::kan* | *BKK12290 → PY79 |
| CG16 | *ndh::kan yfhS::erm* | CG25→ CG2 |
| BLS73 | *amyE::*$P_{hyspank}$*-yjlC-ndh spec* | pLS28 → PY79 |
| CG189 | Δ*yfhS::erm amyE::*$P_{hyspank}$*-yjlC-ndh spec* | pLS28 → CG2 |
| CG135 | *amyE::*$P_{hyspank}$*-yjlC spec* | pLS27 → PY79 |
| CG136 | Δ*yfhS::erm amyE::*$P_{hyspank}$*-yjlC spec* | pLS27 → CG2 |
| CG143 | *amyE::*$P_{hyspank}$*-ndh spec* | pCG23 → PY79 |
| CG144 | Δ*yfhS::erm amyE::*$P_{hyspank}$*-ndh spec* | pCG23→ CG2 |
| CG67 | *rex::erm* | *BKE05970 → PY79 |
| CG69 | *rex::erm amyE::*$P_{hyspank}$*-yjlC-ndh spec* | pLS28 → CG67 |
| CG68 | *rex::erm* Δ*yfhS* | CG67 → Δ*yfhS* (markerless) |
| CG126 | *sacA::*$P_{yjlC}$*-lux* kan | pCG22 → PY79 |
| CG129 | *rex::erm sacA::*$P_{yjlC}$*-lux* kan | CG76→ CG126 |
| CG133 | *yfhS::erm sacA::*$P_{yjlC}$*-lux* kan | CG2 → CG126 |
| CG130 | *yfhS::erm* (LCI #1) *sacA::*$P_{yjlC}$*-lux* kan | CG7 → CG126 |
| CG127 | *sacA::*$P_{yjlC}$*-lux* kan *amyE::*$P_{hyspank}$*-yjlC-ndh spec* | CG189→ CG126 |
| CG145 | *yfhS::erm sacA::*$P_{yjlC}$*-lux* kan *amyE::*$P_{hyspank}$*-yjlC-ndh spec* | CG2→ CG127 |
| CG198 | *yjlC::erm* | *BKE12280 → PY79 |
| CG194 | *yjlC::erm sacA::*$P_{yjlC}$*-lux* kan | CG198 → CG126 |
| CG207 | *yjlC::erm sacA::*$P_{yjlC}$*-lux* kan *amyE::*$P_{hyspank}$*-yjlC spec* | CG135 → CG194 |
| CG204 | *ndh::erm* | *BKE12290 → PY79 |
| CG208 | *ndh::erm sacA::*$P_{yjlC}$*-lux* kan *amyE::*$P_{hyspank}$*-ndh spec* | CG143 → CG204 |
| CG203 | *sacA::*$P_{yjlC}$*-lux* kan *bkdB::Tn917::amyE::*$P_{hyspank}$*-yfhS spec* | CG172 → CG126 |
| CG202 | *yfhS::erm sacA::*$P_{yjlC}$*-lux* kan *bkdB::Tn917::amyE::*$P_{hyspank}$*-yfhS spec* | CG2 → CG203 |
| CG212 | *resD::erm* | *BK323120 → PY79 |
| CG218 | *resD::erm sacA::*$P_{yjlC}$*-lux* kan | *BK323120 → CG126 |
| CG201 | Δ*rex sacA::*$P_{yjlC}$*-lux* kan | CG126 → Δ*rex* markerless |

| | | |
|---|---|---|
| CG213 | *resD::erm* Δ*rex sacA::P$_{yjlC}$-lux* kan | CG212 → CG201 |
| CG221 | *resE::kan* | *BKK23110 → PY79 |
| CG220 | *resE::kan yfhS::erm* | CG221→ CG2 |
| CG186 | *yfhS::*erm *yjlC::*kan | CG2 → CG192 |
| CG192 | *yjlC*::kan | *BKK12280 → PY79 |

### Table S2: Oligonucleotide primers used in this study

| Primer # | Sequence (5'→3') |
|---|---|
| oCG29 | TAGCGGCCGCTGCAGTCC |
| oCG30 | CGAATTCTCATGTTTGACAGCTTATCATCGGC |
| oCG31 | CGATGATAAGCTGTCAAACATGAGAATTCGGAGGAAGCGGTCAAACAG |
| oCG32 | CCCTTTTTTGCCGGACTGCAGCGGCCGCTAATCCAATTCTCCTTTACTATAAGC |
| oCG43 | AATGA TCTAGA TATGCCAGAAACAATCGATC |
| oRB59 | AATAAGTCGACACATAAGGAGGAACTACTATGTATGTCGGACGTGATATGAGCGAA |
| oRB60 | AATAA GCTAGCTTAATCGTAAGAGACGCGCGTGCCGTGGCT |
| oLS41 | AATGAGTCGACACATAAGGAGGAACTACTATGCCAGAAACAATCGATCAAACAAATGCG |
| oLS42 | AATGAGCTAGCTTATTTTTGGTTTTCGCGTTCGTTCATGACTTCAAGG |
| oLS44 | AATGAGCTAGCTTAGTAAGCCAGGCTGAAAAGTCC |
| oCG59 | AATGAGCTAGCTTAGTAAGCCAGGCTGAAAAGTCCTTTAATATG |

```
                                GXGXXG
BsNdh       ------------MSKHIVILGAGYGGVLSALTVRKHYTKEQARVTVVNKYPTHQIITELH     48
CthNdh      -----------MSKPSIVILGAGYGGIVAALGLQKRLNYNEADITLVNKNDYHYITTELH     49
SaNdh       ---------MAQDRKKVLVLGAGYAGLQTVTKLQKAISTEEAEITLINKNEYHYEATWLH     51
MtbNdh      MSPQQEPTAQPPRRHRVVIIGSGFGGLNAAKKLK----RADVDIKLIARTTHHLFQPLLY     56
                            ::::*:*:.*: :.  ::      :. :.:: :   *     *:

BsNdh       RLAAGNVSEKAVAMPLEKLFKGK-DIDLKIAEVSSFSVDKKEVA---LADGSTLTYDALV     104
CthNdh      QPAAGTMHHDQARVGIKELIDEK-KIKFVKDTVVAIDREQQKVT---LQNG-ELHYDYLV     104
SaNdh       EASAGTLNYEDVLYPVESVLKKD-KVNFVQAEVTKIDRDAKKVE---TNQG-IYDFDILV     106
MtbNdh      QVATGIISEGEIAPPTRVVLRKQRNVQVLLGNVTHIDLAGQCVVSELLGHTYQTPYDSLI     116
            . ::* :         . ::  . .:..    *  :.   : *      .     :* *:

BsNdh       VGLGSVTAYFGIPGLEENSMVLKSAADANKVFQHVEDRVR--EYSKTK--NEADATILIG     160
CthNdh      VGLGSEPETFGIEGLREHAFSINSINSVRIIRQHIEYQFA--KFAAEPERT-DYLTIVVG     161
SaNdh       VALGFVSETFGIEGMKDHAFQIENVITARELSRHIEDKFA--NYAASKEKDDNDLSILVG     164
MtbNdh      VAAGAGQSYFGNDHFAEFAPGMKSIDDALELRGRILSAFEQAERSSDPERRAKLLTFTVV     176
            *. *     **   : : :  ::.   .  :  ::   .   : :          :: :

            GXGXXG
BsNdh       GGGLTGVELVGELADIMPNLA--KKYGVDHKEIKLKLVEAGPKILPVLPDDLIERATASL     218
CthNdh      GAGFTGIEFVGELADRMPELC--AEYDVDPKLVRIVNVEAAPTVLPGFDPALVNYAMDVL     219
SaNdh       GAGFTGVEFLGELTDRIPELC--SKYGVDQNKVKITCVEAAPKMLPMFSEELVNHAVSYL     222
MtbNdh      GAGPTGVEMAGQIAELAEHTLKGAFRHIDSTKARVILLDAAPAVLPPMGAKLGQRAAARL     236
            *.* **:*: *::::   .        :* .  ::  ::*.* :** :   * : *   *

BsNdh       EKRGVEFLTGLPVTNV--EGNVI-DLKD-GSKVVANTFVWTGGVQGNPLV----GESGLE     270
CthNdh      GGKGVEFKIGTPIKRCTPEGVVI-EVDGEEEEIKAATVVWTGGVRGNSIL----EKSGFE     274
SaNdh       EDRGVEFKIATPIVACNEKGFVV-EVDGEKQQLNAGTSVWAAGVRGSKLM----EESFEG     277
MtbNdh      QKLGVEIQLGAMVTDVDRNGITVKDSDGTVRRIESACKVWSAGVSASRLGRDLAEQSRVE     296
               ***:  .  :     :* .: : ..   .: :   **:.** .. :      :*

                                                      AQXAXQ
BsNdh       V-NRGRATVNDFLQSTSHEDVFVAGDSAVYFG-PDGRPYPPTAQIAWQMGELIGYNLFAY     328
CthNdh      T-MRGRIKVDPYLRAPGHENIFIVGDCALIINEENNRPYPPTAQIAIQHGENVAANLASL     333
SaNdh       V-KRGRIVTKQDLTINGYDNIFVIGDCSAFIPAGEERPLPTTAQIAMQQGESVAKNIKRI     336
MtbNdh      LDRAGRVQVLPDLSIPGYPNVFVVGDMAAVEG------VPGVAQGAIQGAKYVASTIKAE     350
                **  .   *   .: ::*: ** :           * .** * * .: :. .:

BsNdh       LEGK---TLETFKPVNSGTLASLGRKDAVAIIGANSTPLKGLPASLMKEASNVRYLTHIK     385
CthNdh      IRGG---SMTPFKPHIRGTVASLGRNDAIGIVGG--RKVYGHAASWLKKLIDMRYLYLIG     388
SaNdh       LNGE---STEEFEYVDRGTVCSLGSHDGVGMVFG--KPIAGKKAAFMKKVIDTRAVFKIG     391
MtbNdh      LAGANPAEREPFQYFDKGSMATVSRFSAVAKIGP--VEFSGFIAWLIWLVLHLAYLIGFK     408
            : *        *:    *::.::.  ..:. :      . *  *  :    .   :  :

BsNdh       GL----------FSLAY--------------------------------------          392
CthNdh      GL----------SLVLKKGRF----------------------------------          399
SaNdh       GI----------GLAFKKGKF----------------------------------          402
MtbNdh      TKITTLLSWTVTFLSTRRGQLTITDQQAFARTRLEQLAELAAEAQGSAASAKVAS          463
```

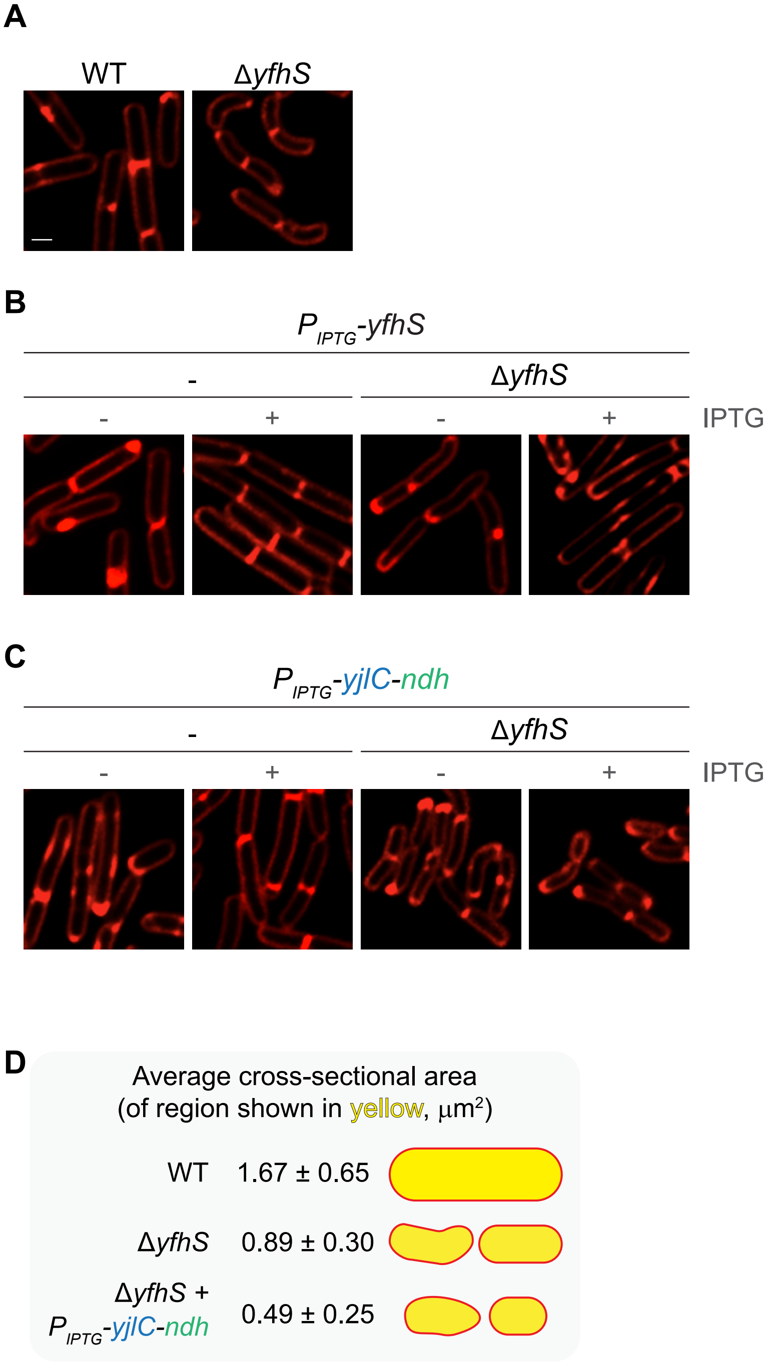
A
WT
ΔyfhS
B
P IPTG-yfhS
-
ΔyfhS
- + - +
IPTG
C
P IPTG-yjlC-ndh
-
ΔyfhS
- + - +
IPTG
D
Average cross-sectional area
(of region shown in yellow, μm²)
WT 1.67 ± 0.65
ΔyfhS 0.89 ± 0.30
ΔyfhS +
P IPTG-yjlC-ndh 0.49 ± 0.25

Figure S2

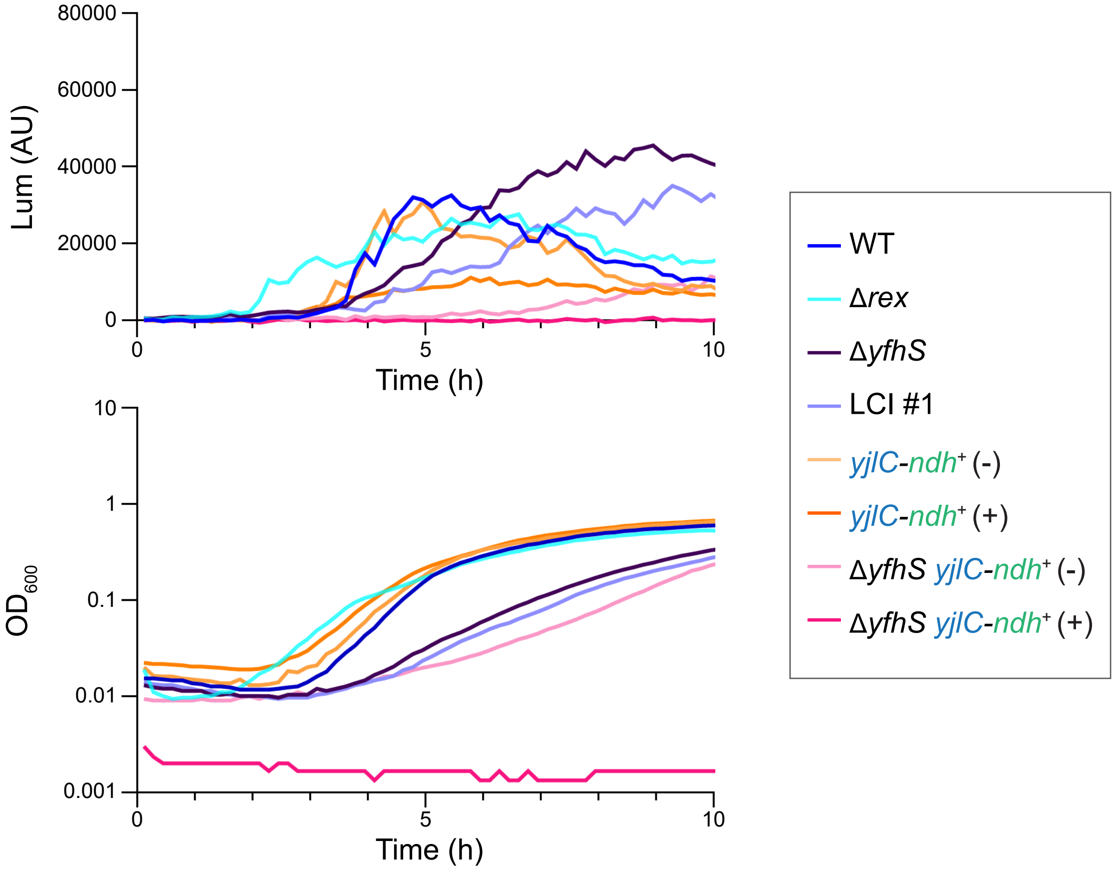

80000
60000
40000
20000
0
Lum (AU)
0
5
10
Time (h)
10
1
0.1
0.01
0.001
$OD_{600}$
0
5
10
Time (h)
WT
Δrex
ΔyfhS
LCI #1
yjlC-ndh⁺ (-)
yjlC-ndh⁺ (+)
ΔyfhS yjlC-ndh⁺ (-)
ΔyfhS yjlC-ndh⁺ (+)


Figure S3

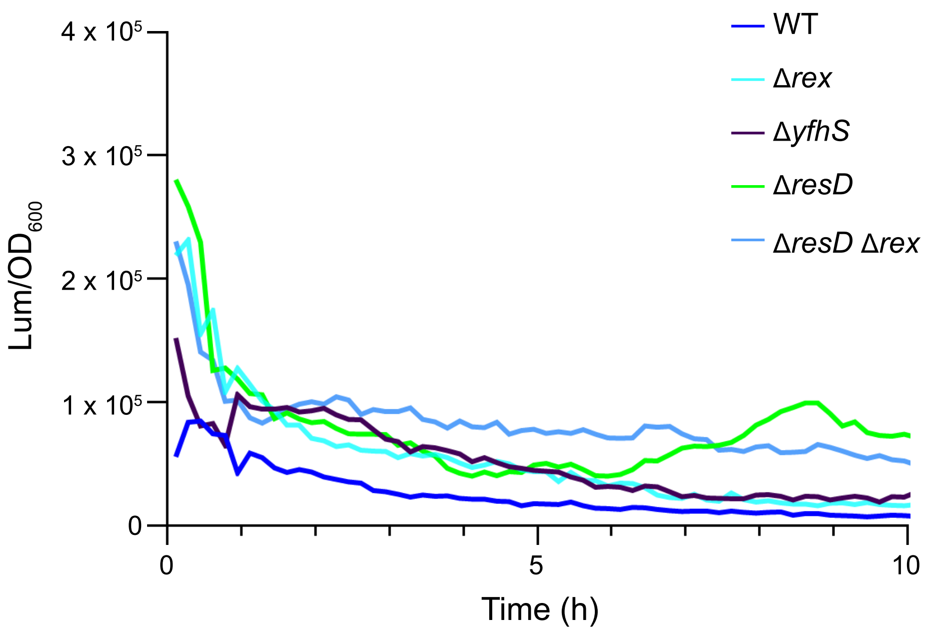

WT
Δrex
ΔyfhS
ΔresD
ΔresD Δrex
Lum/OD600
4 x 10^5
3 x 10^5
2 x 10^5
1 x 10^5
0
0
5
10
Time (h)


Figure S4

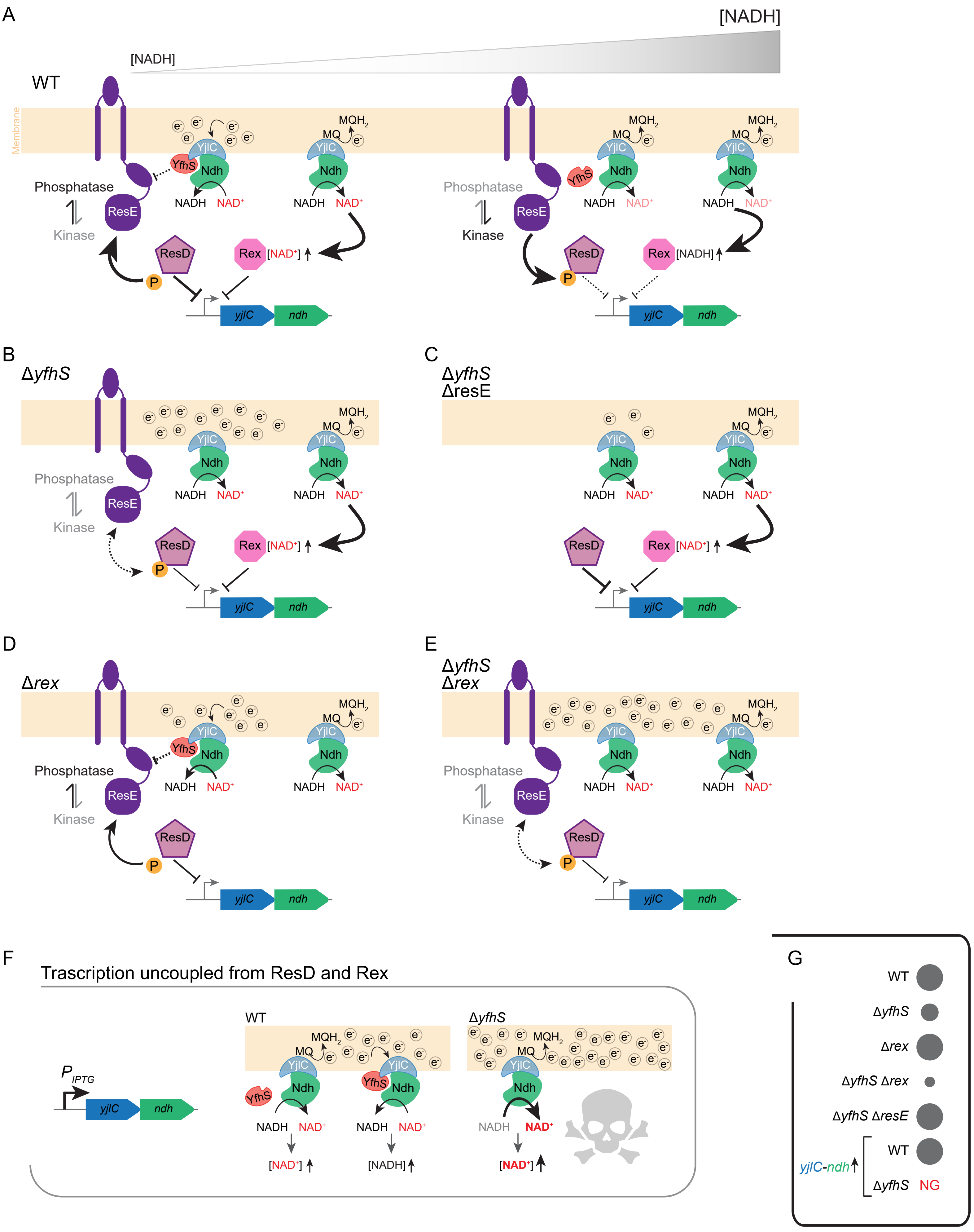
A
[NADH]
[NADH]
WT
Membrane
Phosphatase
Kinase
ResE
YfhS
YjlC
Ndh
NADH
NAD+
MQ
MQH2
ResD
Rex
[NAD+]
[NADH]
P
yjlC
ndh
B
ΔyfhS
C
ΔyfhS
ΔresE
D
Δrex
E
ΔyfhS
Δrex
F
Trascription uncoupled from ResD and Rex
P IPTG
WT
ΔyfhS
G
WT
ΔyfhS
Δrex
ΔyfhS Δrex
ΔyfhS ΔresE
yjlC-ndh
NG

Figure S5

A

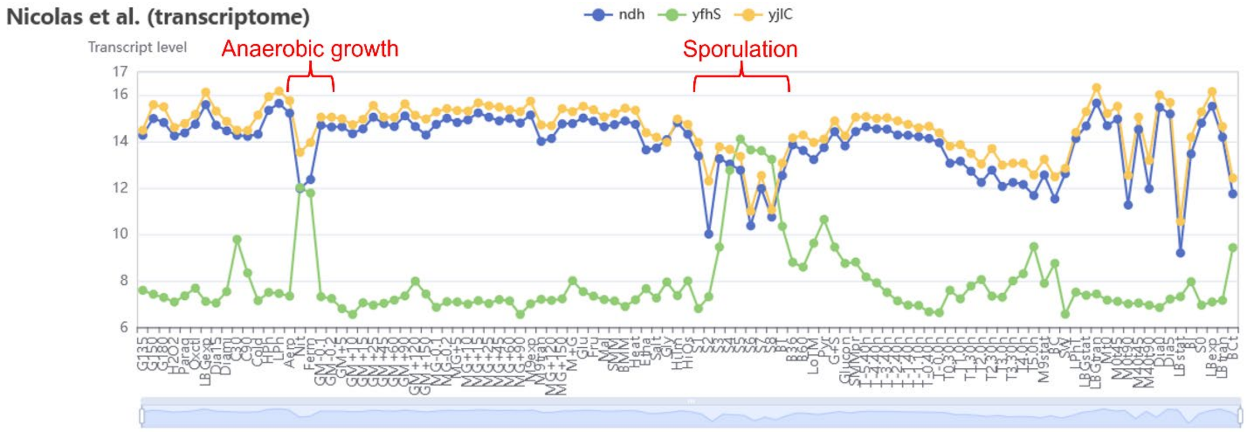
Nicolas et al. (transcriptome)
ndh
yfhS
yjlC
Transcript level
Anaerobic growth
Sporulation
17
16
14
12
10
8
6

B

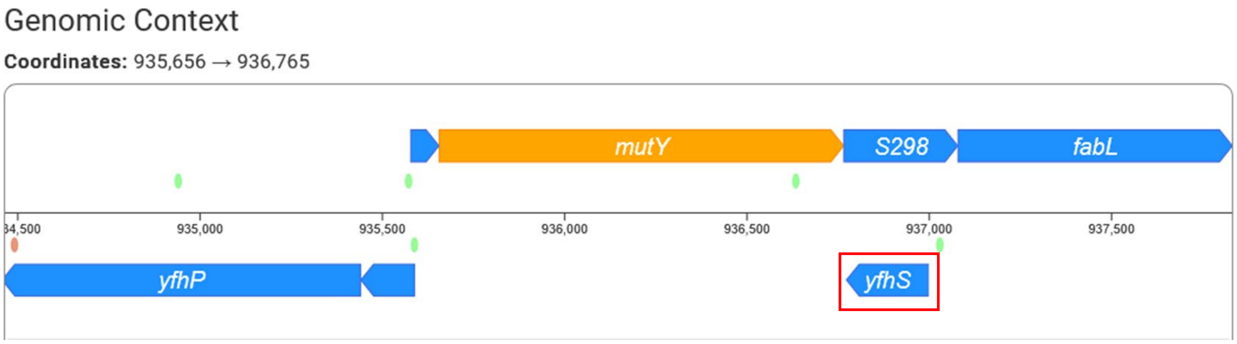
Genomic Context
Coordinates: 935,656 → 936,765
mutY
S298
fabL
935,000
935,500
936,000
936,500
937,000
937,500
yfhP
yfhS

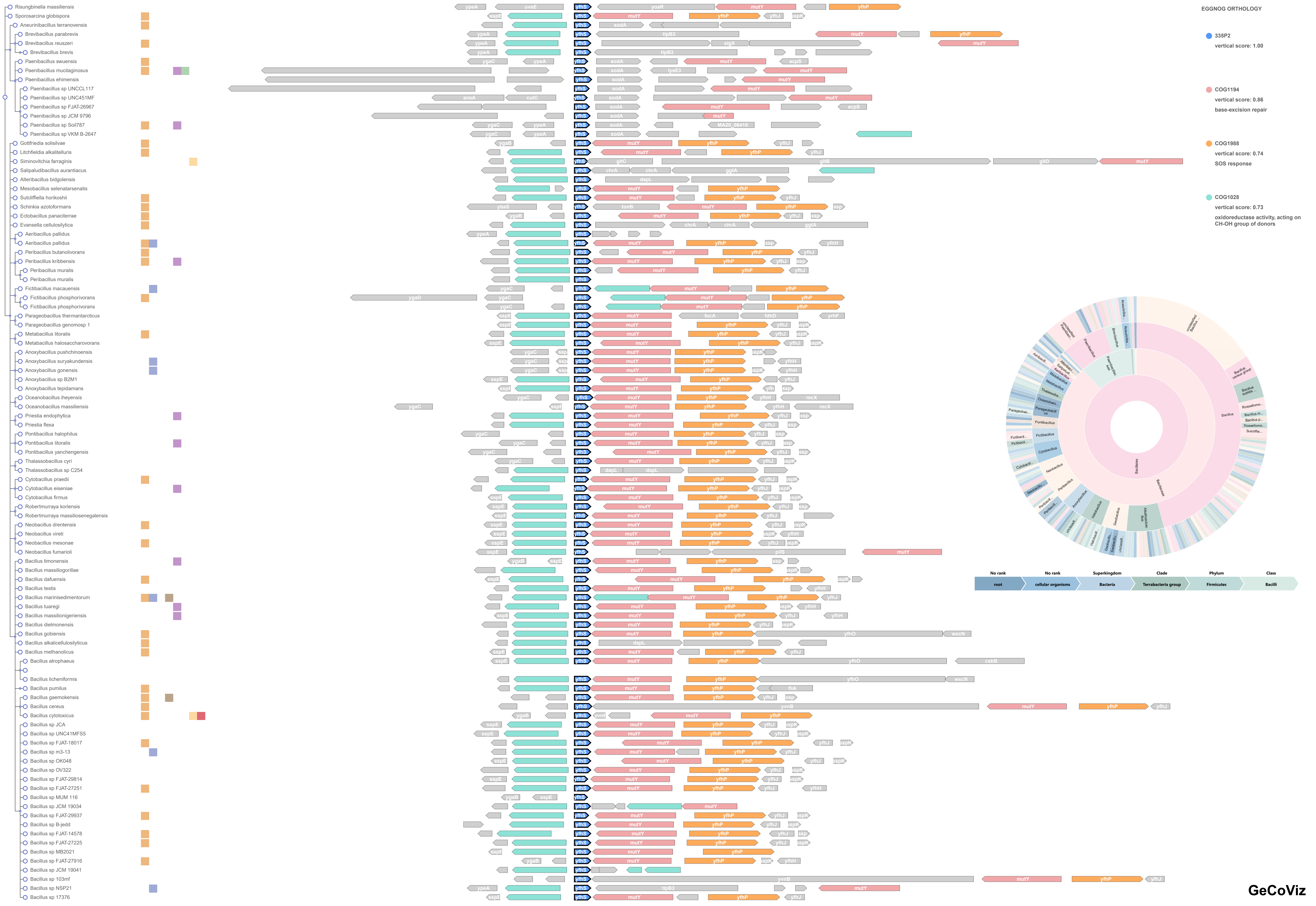
Risungbinella massiliensis
Sporosarcina globispora
Aneurinibacillus terranovensis
Brevibacillus parabrevis
Brevibacillus reuszeri
Brevibacillus brevis
Paenibacillus swuensis
Paenibacillus mucilaginosus
Paenibacillus ehimensis
Paenibacillus sp UNCCL117
Paenibacillus sp UNC451MF
Paenibacillus sp FJAT-26967
Paenibacillus sp JCM 9796
Paenibacillus sp Soil787
Paenibacillus sp VKM B-2647
Gottfriedia solisilvae
Litchfieldia alkalitelluris
Siminovitchia farraginis
Salipaludibacillus aurantiacus
Alteribacillus bidgolensis
Mesobacillus selenatarsenatis
Sutcliffiella horikoshii
Schinkia azotoformans
Ectobacillus panaciterrae
Evansella cellulosilytica
Aeribacillus pallidus
Aeribacillus pallidus
Peribacillus butanolivorans
Peribacillus kribbensis
Peribacillus muralis
Peribacillus muralis
Fictibacillus macauensis
Fictibacillus phosphorivorans
Fictibacillus phosphorivorans
Parageobacillus thermantarcticus
Parageobacillus genomosp 1
Metabacillus litoralis
Metabacillus halosaccharovorans
Anoxybacillus pushchinoensis
Anoxybacillus suryakundensis
Anoxybacillus gonensis
Anoxybacillus sp B2M1
Anoxybacillus tepidamans
Oceanobacillus iheyensis
Oceanobacillus massiliensis
Priestia endophytica
Priestia flexa
Pontibacillus halophilus
Pontibacillus litoralis
Pontibacillus yanchengensis
Thalassobacillus cyri
Thalassobacillus sp C254
Cytobacillus praedii
Cytobacillus eiseniae
Cytobacillus firmus
Robertmurraya korlensis
Robertmurraya massiliosenegalensis
Neobacillus drentensis
Neobacillus vireti
Neobacillus mesonae
Neobacillus fumarioli
Bacillus timonensis
Bacillus massiliogorillae
Bacillus dafuensis
Bacillus testis
Bacillus marinisedimentorum
Bacillus tuaregi
Bacillus massilionigeriensis
Bacillus dielmonensis
Bacillus gobiensis
Bacillus alkalicellulosilyticus
Bacillus methanolicus
Bacillus atrophaeus
Bacillus licheniformis
Bacillus pumilus
Bacillus gaemokensis
Bacillus cereus
Bacillus cytotoxicus
Bacillus sp JCA
Bacillus sp UNC41MFS5
Bacillus sp FJAT-18017
Bacillus sp m3-13
Bacillus sp OK048
Bacillus sp OV322
Bacillus sp FJAT-29814
Bacillus sp FJAT-27251
Bacillus sp MUM 116
Bacillus sp JCM 19034
Bacillus sp FJAT-29937
Bacillus sp B-jedd
Bacillus sp FJAT-14578
Bacillus sp FJAT-27225
Bacillus sp MB2021
Bacillus sp FJAT-27916
Bacillus sp JCM 19041
Bacillus sp 103mf
Bacillus sp NSP21
Bacillus sp 17376
EGGNOG ORTHOLOGY
335P2
vertical score: 1.00
COG1194
vertical score: 0.86
base-excision repair
COG1988
vertical score: 0.74
SOS response
COG1028
vertical score: 0.73
oxidoreductase activity, acting on CH-OH group of donors
No rank
root
No rank
cellular organisms
Superkingdom
Bacteria
Clade
Terrabacteria group
Phylum
Firmicutes
Class
Bacilli
GeCoViz

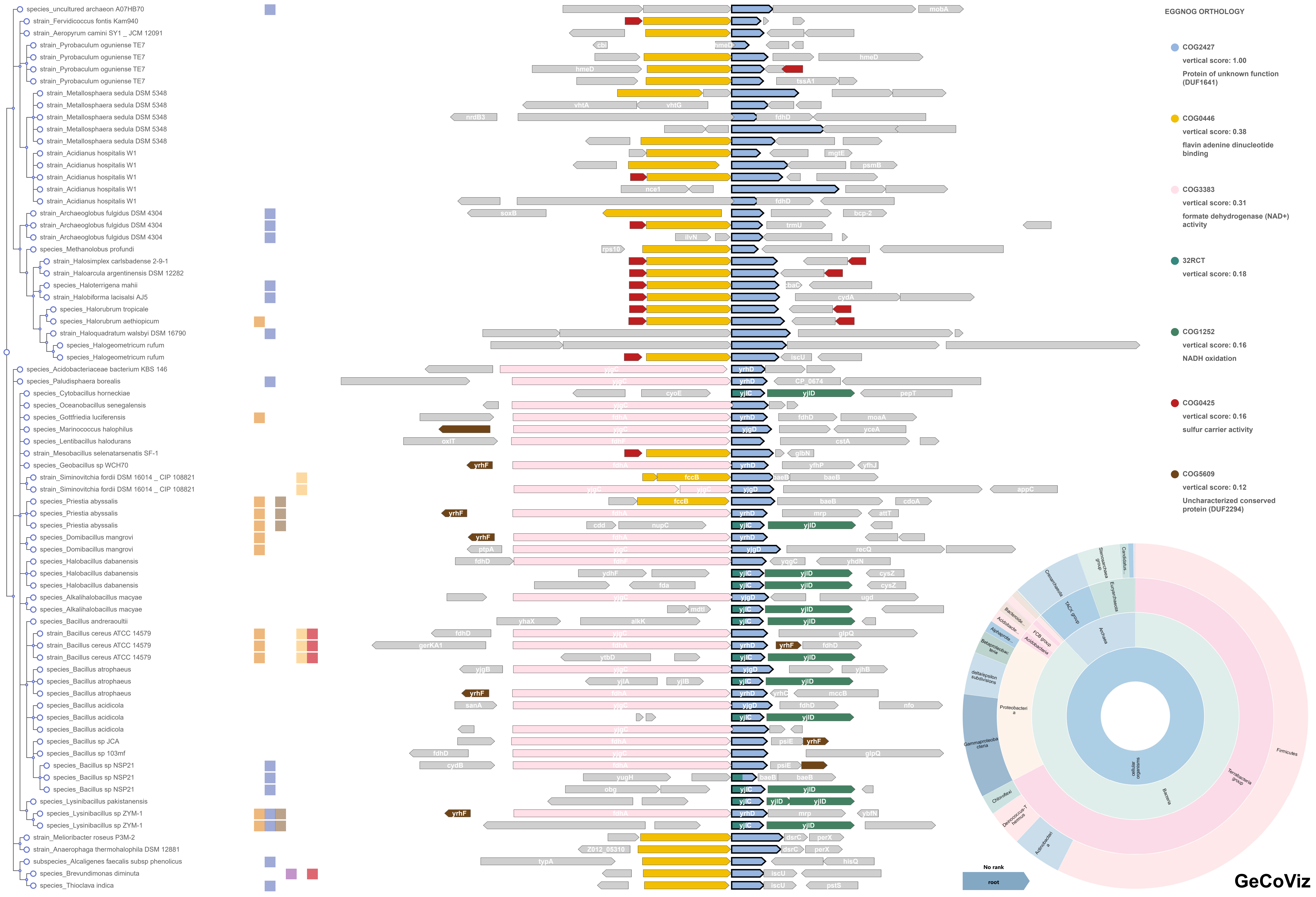
species_uncultured archaeon A07HB70
strain_Fervidicoccus fontis Kam940
strain_Aeropyrum camini SY1 _ JCM 12091
strain_Pyrobaculum oguniense TE7
strain_Pyrobaculum oguniense TE7
strain_Pyrobaculum oguniense TE7
strain_Pyrobaculum oguniense TE7
strain_Metallosphaera sedula DSM 5348
strain_Metallosphaera sedula DSM 5348
strain_Metallosphaera sedula DSM 5348
strain_Metallosphaera sedula DSM 5348
strain_Metallosphaera sedula DSM 5348
strain_Acidianus hospitalis W1
strain_Acidianus hospitalis W1
strain_Acidianus hospitalis W1
strain_Acidianus hospitalis W1
strain_Acidianus hospitalis W1
strain_Archaeoglobus fulgidus DSM 4304
strain_Archaeoglobus fulgidus DSM 4304
strain_Archaeoglobus fulgidus DSM 4304
species_Methanolobus profundi
strain_Halosimplex carlsbadense 2-9-1
strain_Haloarcula argentinensis DSM 12282
species_Haloterrigena mahii
strain_Halobiforma lacisalsi AJ5
species_Halorubrum tropicale
species_Halorubrum aethiopicum
strain_Haloquadratum walsbyi DSM 16790
species_Halogeometricum rufum
species_Halogeometricum rufum
species_Acidobacteriaceae bacterium KBS 146
species_Paludisphaera borealis
species_Cytobacillus horneckiae
species_Oceanobacillus senegalensis
species_Gottfriedia luciferensis
species_Marinococcus halophilus
species_Lentibacillus halodurans
strain_Mesobacillus selenatarsenatis SF-1
species_Geobacillus sp WCH70
strain_Siminovitchia fordii DSM 16014 _ CIP 108821
strain_Siminovitchia fordii DSM 16014 _ CIP 108821
species_Priestia abyssalis
species_Priestia abyssalis
species_Priestia abyssalis
species_Domibacillus mangrovi
species_Domibacillus mangrovi
species_Halobacillus dabanensis
species_Halobacillus dabanensis
species_Halobacillus dabanensis
species_Alkalihalobacillus macyae
species_Alkalihalobacillus macyae
species_Bacillus andreraoultii
strain_Bacillus cereus ATCC 14579
strain_Bacillus cereus ATCC 14579
strain_Bacillus cereus ATCC 14579
species_Bacillus atrophaeus
species_Bacillus atrophaeus
species_Bacillus atrophaeus
species_Bacillus acidicola
species_Bacillus acidicola
species_Bacillus acidicola
species_Bacillus sp JCA
species_Bacillus sp 103mf
species_Bacillus sp NSP21
species_Bacillus sp NSP21
species_Bacillus sp NSP21
species_Lysinibacillus pakistanensis
species_Lysinibacillus sp ZYM-1
species_Lysinibacillus sp ZYM-1
strain_Melioribacter roseus P3M-2
strain_Anaerophaga thermohalophila DSM 12881
subspecies_Alcaligenes faecalis subsp phenolicus
species_Brevundimonas diminuta
species_Thioclava indica
EGGNOG ORTHOLOGY
COG2427
vertical score: 1.00
Protein of unknown function (DUF1641)
COG0446
vertical score: 0.38
flavin adenine dinucleotide binding
COG3383
vertical score: 0.31
formate dehydrogenase (NAD+) activity
32RCT
vertical score: 0.18
COG1252
vertical score: 0.16
NADH oxidation
COG0425
vertical score: 0.16
sulfur carrier activity
COG5609
vertical score: 0.12
Uncharacterized conserved protein (DUF2294)
cellular organisms
Bacteria
Archaea
Firmicutes
Terrabacteria group
Proteobacteria
Gammaproteobacteria
Chloroflexi
Actinobacteria
Crenarchaeota
TACK group
Euryarchaeota
Stenosarchaea group
Acidobacteria
FCB group
No rank
root
GeCoViz
